# Single-cell transcriptomics reveal distinctive patterns of fibroblast activation in murine heart failure with preserved ejection fraction

**DOI:** 10.1101/2023.05.09.539983

**Authors:** Jan D. Lanzer, Laura M. Wienecke, Ricardo O. Ramirez-Flores, Maura M. Zylla, Niklas Hartmann, Florian Sicklinger, Jobst-Hendrick Schultz, Norbert Frey, Julio Saez-Rodriguez, Florian Leuschner

## Abstract

Inflammation, fibrosis and metabolic stress critically promote heart failure with preserved ejection fraction (HFpEF). Exposure to high-fat diet and nitric oxide synthase inhibitor N[w]-nitro-l-arginine methyl ester (L-NAME) recapitulate features of HFpEF in mice. To identify disease specific traits during adverse remodeling, we profiled interstitial cells in early murine HFpEF using single-cell RNAseq (scRNAseq). Diastolic dysfunction and perivascular fibrosis were accompanied by an activation of cardiac fibroblast and macrophage subsets. Integration of fibroblasts from HFpEF with two murine models for heart failure with reduced ejection fraction (HFrEF) identified a catalog of conserved fibroblast phenotypes across mouse models. Moreover, HFpEF specific characteristics included induced metabolic, hypoxic and inflammatory transcription factors and pathways, including enhanced expression of Angiopoietin-like 4 next to basement membrane compounds. Fibroblast activation was further dissected into transcriptional and compositional shifts and thereby highly responsive cell states for each HF model were identified. In contrast to HFrEF, where myofibroblast and matrifibrocyte activation were crucial features, we found that these cell-states played a subsidiary role in early HFpEF. These disease-specific fibroblast signatures were corroborated in human myocardial bulk transcriptomes. Furthermore, we found an expansion of pro-inflammatory Ly6C^high^ macrophages in HFpEF, and we identified a potential cross-talk between macrophages and fibroblasts via SPP1 and TNFɑ. Finally, a marker of murine HFpEF fibroblast activation, Angiopoietin-like 4, was elevated in plasma samples of HFpEF patients and associated with disease severity. Taken together, our study provides a comprehensive characterization of molecular fibroblast and macrophage activation patterns in murine HFpEF, as well as the identification of a novel biomarker for disease progression in patients.

## Introduction

Heart failure with preserved ejection fraction (HFpEF) represents one of the largest unmet clinical needs in cardiovascular medicine, given that it accounts for about 50% of heart failure (HF) patients and is increasing in prevalence^1^. However, apart from gliflozins, no effective treatment strategies exist to reduce the associated diastolic dysfunction, fibrosis, hypertrophy and the resulting pronounced morbidity and mortality. Therapeutic concepts and established drugs for the treatment of heart failure with reduced ejection fraction (HFrEF) failed broadly when tested for beneficial effects in HFpEF, suggesting fundamentally different pathomechanisms^1^.

HFpEF comprises a complex and multifactorial interplay of the disease promoting risk factors, such as hypertension, obesity, metabolic syndrome, chronic inflammation, and aging. Suitable animal models were missing until a few years ago, when a two-hit mouse model combining a 60% high-fat diet with inhibition of the constitutive nitric oxide synthase by Nω-nitro-l-arginine methyl ester (L-NAME) recapitulated metabolic and hypertensive stress in HFpEF^2, 3^. Analysis of this model led to major mechanistic insights in the pathophysiology of hypertrophy and cardiac immunometabolic alterations in HFpEF^3–5^ and potential therapeutic targets. Since these studies focused predominantly on cardiomyocyte hypertrophy and metabolism^1^, little knowledge was gathered about the distinct role of cardiac interstitial cells and their cross-talk in ventricular stiffening and fibrosis^1, 4^.

Single-cell RNA sequencing (scRNAseq) allows for the quantification of transcriptional changes of individual cells and description of cell phenotype heterogeneity. Consequently, scRNAseq has opened the door for fundamental insights into cellular heterogeneity, developmental biology and molecular disease processes in the cardiovascular field^6–8^. Thus, its application to a HFpEF model could shed light on the cellular disease mechanisms.

Here we present to our knowledge the first scRNAseq analysis of the ventricular interstitium in mice receiving L-NAME and high fat diet (further called HFpEF model) in early stages of diastolic dysfunction. We compared fibroblast phenotypes and disease signatures by integration with scRNAseq data from other HF models that recapitulate HFrEF and identified HFpEF specific patterns of fibroblast activation. We characterized HFpEF associated fibrotic signatures and compared them with human bulk references, providing new pathophysiological hypotheses relevant for the understanding of fibrosis in HFpEF necessary for future anti-fibrotic drug development.

## Results

### Disease model and data description

To mimic HFpEF, we used the established two-hit mouse model that induces metabolic and hypertensive stress by 60% high-fat diet and L-NAME, respectively^3^. From seven weeks of dietary intervention onwards, a diastolic dysfunction phenotype was observed echocardiographically under preservation of systolic left ventricular function (Fig. 1A, Supp. Fig. 1). Body and heart weight, normalized to tibia length, increased concordantly indicating obesity and cardiac hypertrophy (Fig. 1A, Supp. Fig. 1A-C). To describe this early remodeling, we isolated cardiac interstitial cells after seven weeks by MACS^Ⓡ^ dead cell depletion and FACS sorting of live and metabolically active cells (Fig. 1B). We performed scRNAseq with the 10x Chromium droplet based platform to analyze cellular transcriptomic changes within cardiac ventricular interstitial cells of two control and two HFpEF murine hearts. After processing and quality control we retained expression profiles of 6,132 cells described by 15,046 genes (mean UMI coverage per cell: 2,838) (Supp. Fig. 2). Unsupervised clustering yielded 10 distinct clusters (Fig. 1C) representing major cell types of the cardiac interstitium based on their top marker genes and known canonical markers. We identified two fibroblast clusters (Col1a1+ and Wif1+), endothelial cells (EC) (Pecam1+), natural killer cells (Gzma+), macrophages (CD68+), T effector cells (CD8+) and T helper cells (CD4+), B cells (CD19+), granulocytes (S100a9+), smooth muscle cells and pericytes (Acta2+) (Figure 1D).

**Figure 1.**
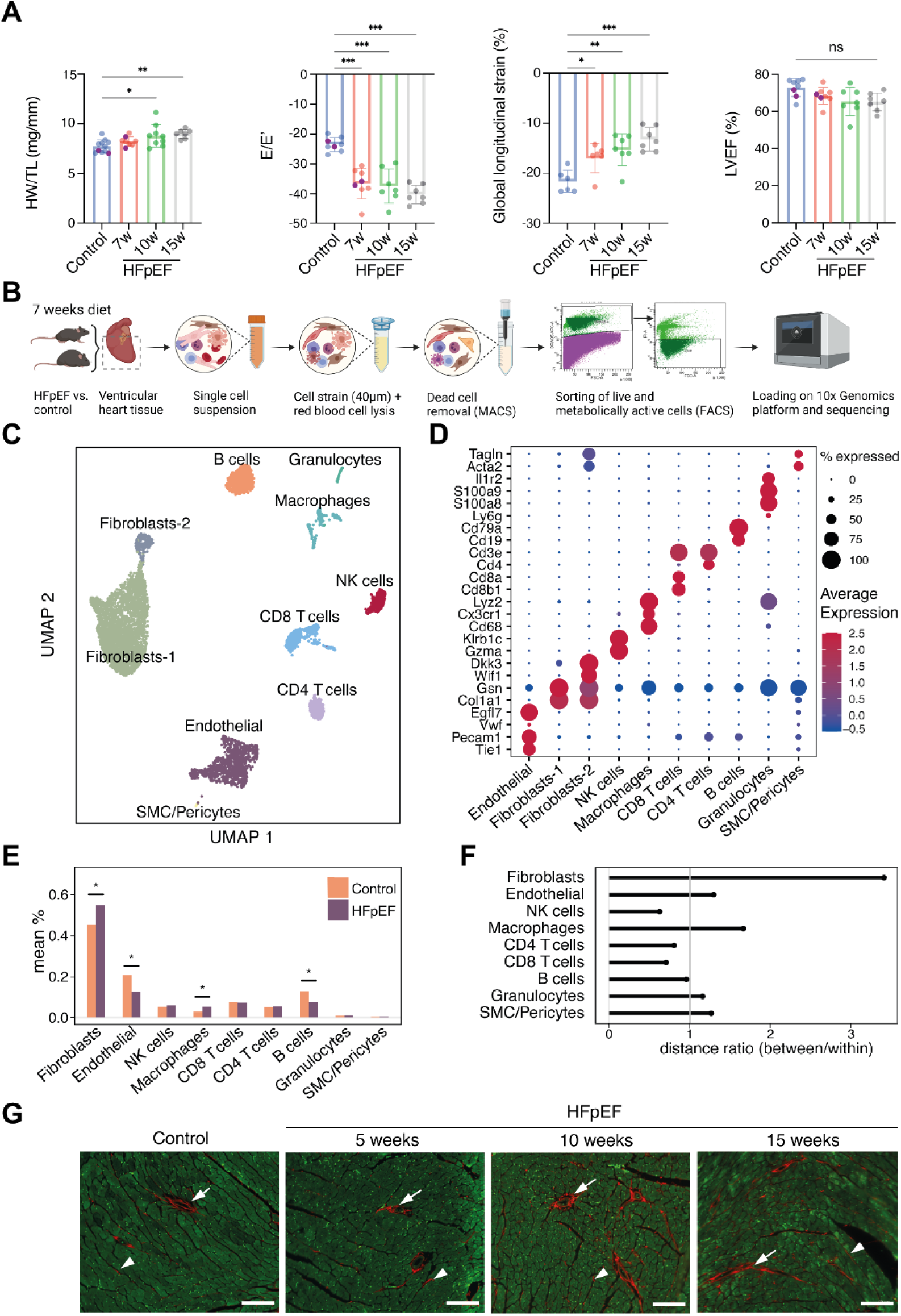
Study model and cell type assignment. A) Murine HFpEF model characterization by ratio of heart weight to tibia length (HW/TL) and echocardiographic hallmarks (E/E’, global-longitudinal strain and LVEF), purple data points represent the animals used for single-cell RNA sequencing (scRNAseq). Statistical analysis performed by one-way ANOVA, bar graphs indicate mean±SD, *p<0.05, **p<0.01, ***p<0.001. ns= deemed not significant (p>0.05), LVEF= left ventricular ejection fraction, w= weeks. B) Schematic summary of experimental setup for scRNAseq experiments using mice after 7 weeks of HFpEF or control diet. C) UMAP embeddings of normalized scRNAseq data after processing and filtering. D) Marker gene expression for cell type assignment. E) Cell type composition of main cell types as mean percentage per group, compared between HFpEF and control mice. *p < 0.05, p-values were calculated via label permutation. F) Cosine distance ratios of highly variable genes between pseudobulked cell type profiles. Median between group distance is divided by median within group distance. G) Representative Picrosirius-Red stainings of interstitial fibrotic fibers (arrow heads) and perivascular fibrosis (arrows) from control and different stages of HFpEF heart sections. Imaging performed in 594nm (Picrosirius-Red) and 488nm (autofluorescence) channels. White scale bars in the right bottom corner correspond to 100μm.

**Figure 2.**
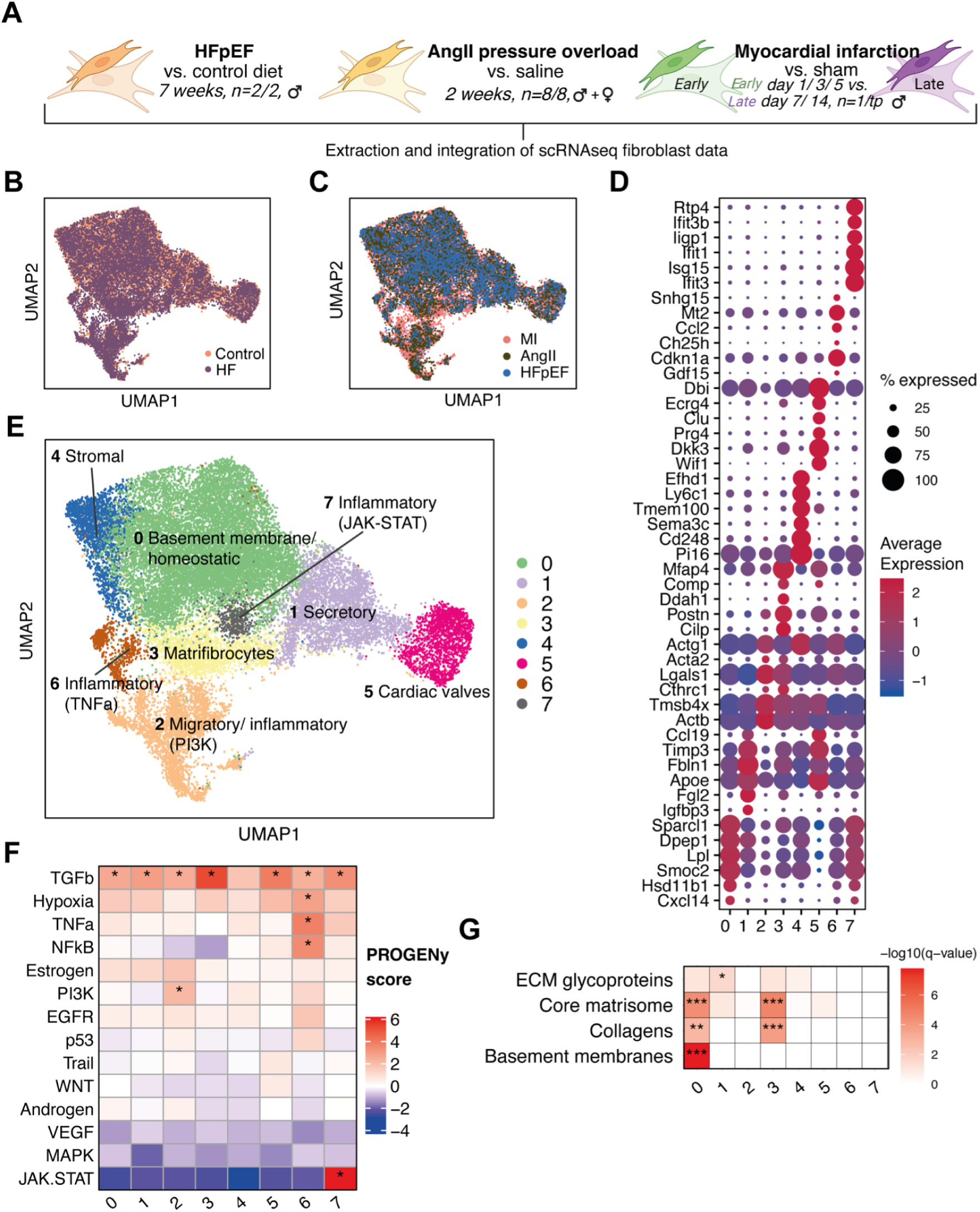
Integrated atlas of cardiac fibroblasts from different disease models. A) Schematic of the integrated murine HFpEF and HFrEF (AngII and MI) fibroblast studies. B+C) UMAP embeddings of integrated fibroblasts, colored by disease (HF, Heart Failure) vs. control (B), study (C). D) Overview of top cell state marker expression of integrated fibroblast states. E) UMAP embeddings, showing the integrated fibroblast atlas colored by cell clusters, i.e. the integrated fibroblast states (IFS). Labels indicate possible fibroblast differentiations based on functional characterization. F) Estimated pathway activities with PROGENy based on effect size (avg log2 fold change) of footprint genes in integrated fibroblast states. *PROGENy z-score > 2. G) Overrepresentation analysis of extracellular matrix related gene sets with markers of integrated fibroblast states. Hypergeometric test with Benjamini Hochberg correction, *q < 0.05, **q < 0.01, ***q < 0.001.

### Cell type composition and molecular profiles suggest fibroblast and macrophage involvement in cardiac remodeling

To identify interstitial cells involved in HFpEF remodeling, we first compared the cellular composition in control with HFpEF cardiac tissue, and evaluated the significance of compositional changes by label permutation (see methods). This yielded a modest increase of fibroblasts and macrophages and decrease of B cells and ECs in HFpEF (Fig. 1E, Supp. Fig. 3A-B).

**Figure 3.**
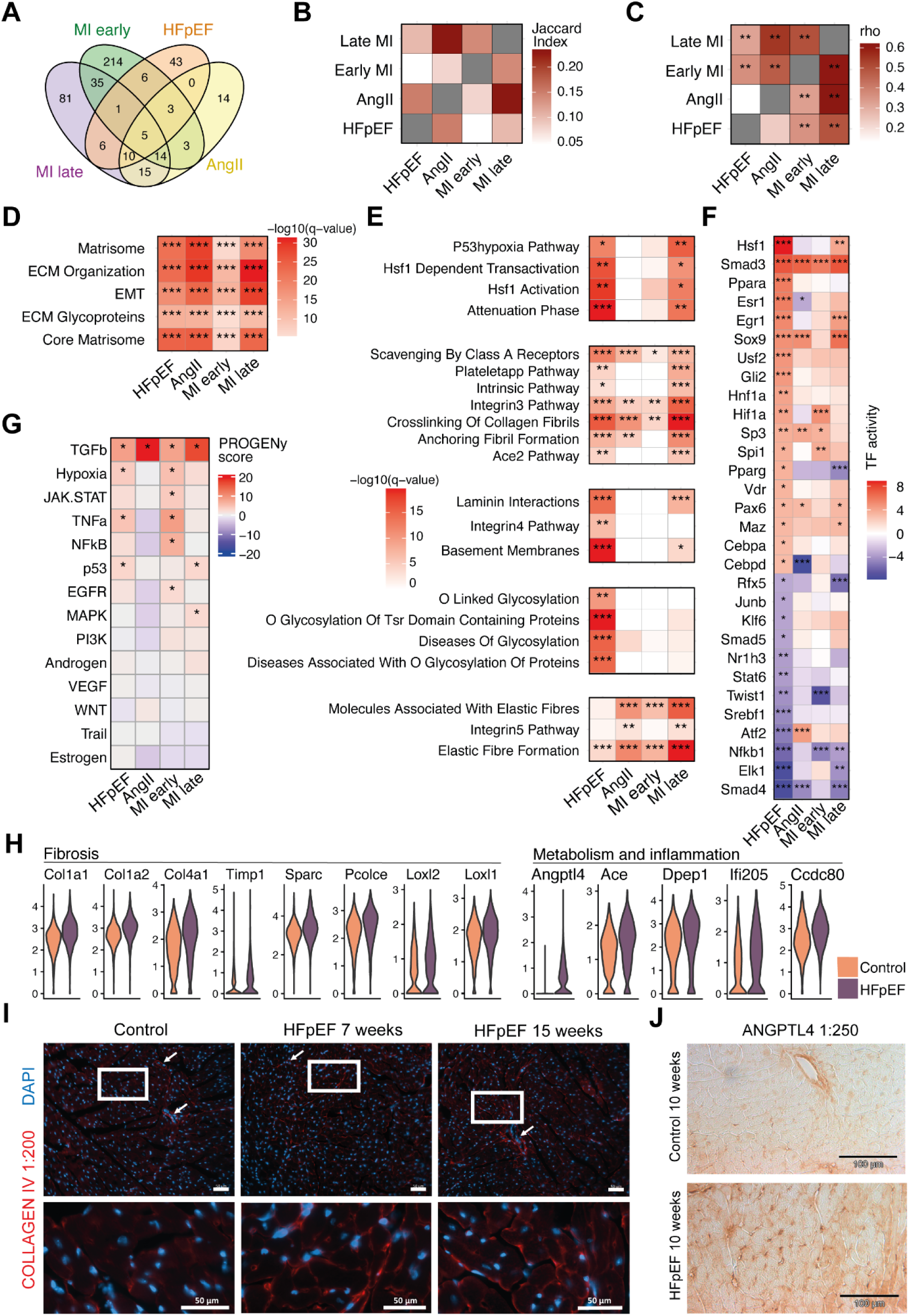
Comparison and interpretation of fibroblast disease signatures from different heart failure models. A) Comparing intersections of upregulated genes in different heart failure (HF) models. B) Intersection quantification via jaccard index. C) Comparison of direction of regulation between studies. Pearson correlation was calculated between log fold change vectors of signature genes in pairwise comparisons. Each study comparison was based on the upregulated genes from the study on the x-axis. **p < 0.01. D+E) Heatmaps of geneset overrepresentation in study specific fibroblast disease signatures. Significantly enriched gene sets common between signatures (D) and selected gene sets to highlight HFpEF signature characteristics (E), ECM = extracellular matrix, EMT = epithelial-mesenchymal transition. Each heatmap in E) represents a group of similar genesets. Hypergeometric test with Benjamini Hochberg correction, *q <0.01, **q<0.001, ***q<0.0001. F) Estimated transcription factor activities with DoRothEA based on effect size (log fold change) of target genes compared between HF models. G) Estimated pathway activities with PROGENy based on effect size (log fold change) of footprint genes compared between HF models. H) Expression values of selected fibrosis and inflammatory genes in individual fibroblasts in HFpEF (grey) and control (orange) mice. All genes were significantly upregulated (adj. p-value < 0.05). I) Immunofluorescence images of collagen IV (red) and DAPI (blue) staining of left ventricular heart sections. Lower panels show magnifications of the areas marked by white boxes. White arrows indicate capillaries or larger blood vessels. Scale bars in the right bottom corner indicate 50 μm length. J) Immunohistological staining of Angptl4 protein in left ventricular heart sections.

As cell type compositions are not independent and therefore only partially informative of the importance of a cell type for disease process, we assessed whether the variation of gene expression between experimental groups was higher than the variability expected within a single group (see methods) (Fig. 1F). We found that fibroblasts displayed the highest ratio of ‘between to within group distance’ followed by macrophages. ECs and B cells did not display high disease associated variability, suggesting that their relative decrease in proportion is not associated with fundamental gene expression changes. We applied a cell type prioritization method to rank cell types by classifier performance. This classifier was trained to separate healthy from diseased cells and can provide an additional estimate for magnitude of molecular changes in cell types^9^. This yielded the highest performance for macrophages and endothelial cells, followed by modest performance for fibroblasts (Supp. Fig. 3C). L-NAME treatment directly targets ECs, expected to induce direct transcriptional changes. Taken togther, the compositional change and molecular differences suggested that fibroblasts and macrophages are important contributors to the early HFpEF associated remodeling and phenotype.

Fibroblast activity relates to cardiac fibrosis, which is a hallmark feature of human HFpEF^10^. In parallel, we found a qualitative increase of interstitial and perivascular collagen deposition with time in the HFpEF model (Fig. 1G). Thus, seven weeks of HFpEF diet already recapitulated all features of HFpEF including cardiac fibrosis and mild functional changes of the left ventricle at this time-point. While fibrosis represents one of the major pathomechanisms without current mitigating therapeutic options, investigating early fibroblast activation is of high interest to understand HFpEF-related cardiac fibrosis.

### Fibroblast phenotype definitions across murine heart failure models

Cardiac fibroblasts accomplish a wide range of biological functions, crucial for tissue homeostasis and architecture^11^. In human HFrEF and HFpEF, cardiac fibrosis represents a major axis of reparative and adverse remodeling. While histologically HFpEF has been associated with interstitial and perivascular fibrosis, the underlying functional characteristics of fibroblast activation remain unknown^10, 12^. Thus, we sought to compare HFpEF fibroblast activation with other cardiac fibrotic disease etiologies by integration of our single-cell data with two other single-cell resources that represent different types of HFrEF: first, a model for cardiac fibrosis and hypertrophy by hypertensive stress induced by two weeks of angiotensin II (AngII) administration^13^ and second, an acute myocardial infarction (MI) model^7^ that assessed early (<7 days) and later ischemic remodeling (7-14 days) (Fig. 2A). The MI model is characterized by cell death and associated replacement fibrosis^14^ while the AngII administration causes initially extensive reactive fibrosis^15^.

We uniformly processed studies, annotated cell types and identified fibroblasts by selecting Col1a1+, Pdgfra+ and Gsn+ cells (Supp. Fig. 4). Fibroblasts from three datasets were then integrated with *Harmony*^16^ while accounting for sample and study batch effects that resulted in an integrated cardiac fibroblast atlas of 26,455 cells, capturing a wide spectrum of phenotype diversity across HF models. Study and sample batch effects were satisfactorily mitigated (see methods) (Fig. 2B, C).

**Figure 4.**
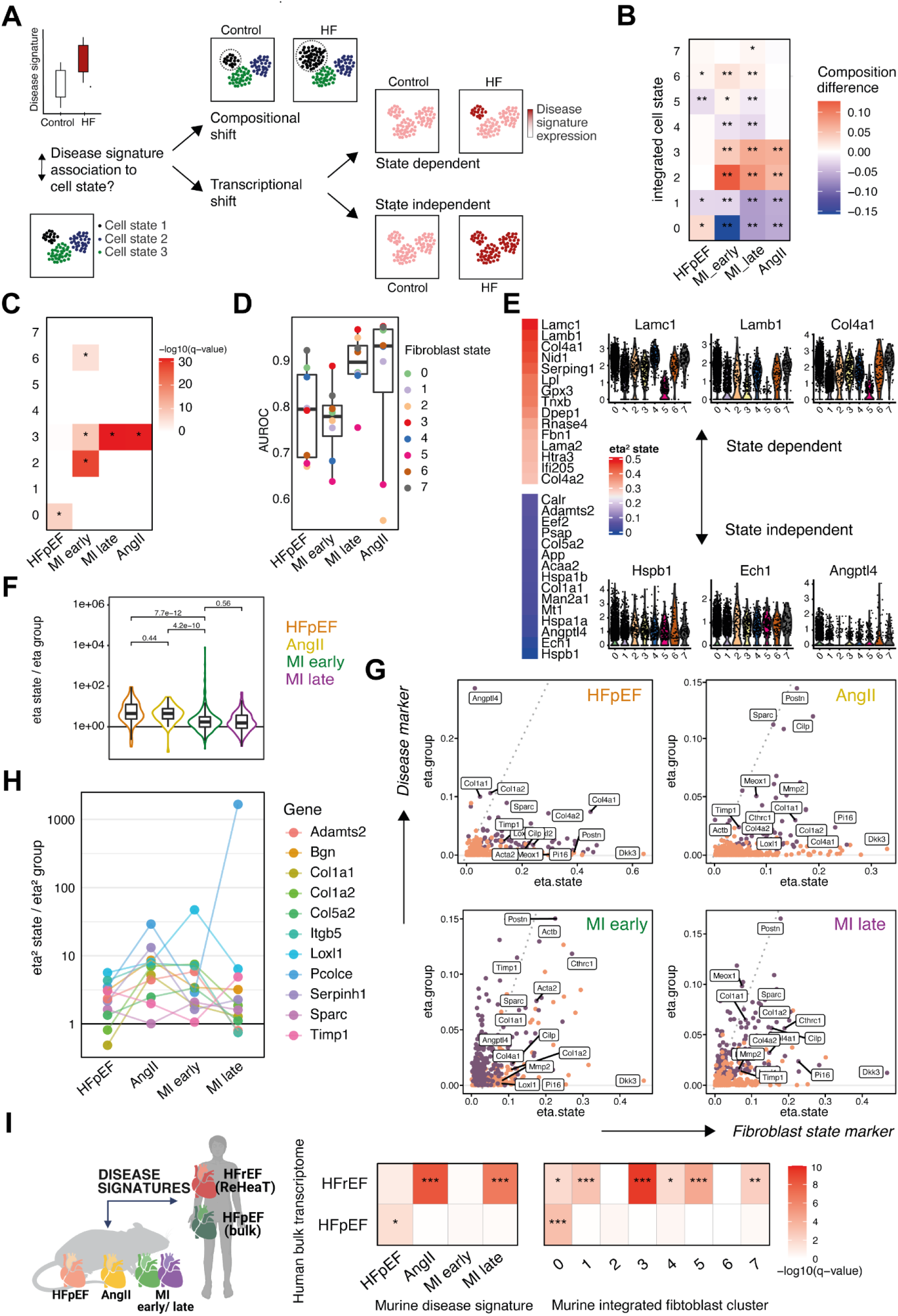
Decomposing Fibroblast disease signatures. A) Schematic of different expression patterns in regard to cell states that could yield an upregulation of a disease signature. Compositional shifts by expanding cell number are distinguished from transcriptional shifts via uniform (state independent) or non-uniform (state dependent) upregulation of disease signatures. B) Composition change of integrated fibroblast states (IFS) between control and heart failure per study. p-values calculated via label permutation, *p < 0.05, **p < 0.01. C) Overrepresentation analysis of disease specific fibroblast signatures (x-axis) and top 100 IFS markers (y-axis). Hypergeometric test, *p < 0.05. D) Gene set scores of study specific signatures (x-axis) were used to calculate the area under the receiver operator curve (AUROC, y-axis) between control and diseased cells within each IFS (color). E) HFpEF signature expression dependent on IFS category by calculating the explained variance (eta² values) of gene-wise ANOVAs. Violin plots display normalized expression values of three genes with lowest (lower panel) and highest (upper panel) variance explained by cell state. F) Quantification of differences in state dependent regulation of disease signatures across heart failure models. The ratio of the explained variance by IFS and disease class was calculated for each HF model and its disease signature. Wilcoxon-test p-values are shown. G) Explained variance (eta² values) by IFS on x-axis and explained variance by disease class (gene ∼ disease class) on y-axis. Violett dots are part of the disease signature. H) The ratio of explained variance by state and disease class of selected genes that were upregulated in all HF models. I) Corroboration of murine fibroblast signatures in human myocardial samples. Human HFpEF and HFrEF studies were curated and top differentially upregulated genes were selected (y-axis). Gene set overlaps with fibroblast disease signatures from different study models (left-panel) or fibroblast state marker (right panel) (hypergeometric test). AngII= angiotensin II model, HFpEF= heart failure with preserved ejection fraction, MI= myocardial infarction. q-value = Benjamini Hochberg corrected p-value, *q <0.05, **q<0.01, ***<0.001.

Previous studies identified cardiac fibroblast phenotypes at the single-cell level in healthy and diseased hearts with limited consistency^7, 13, 17–20^. Thus, the integration allowed us to robustly define high-level fibroblast phenotypes across different cardiac remodeling scenarios that enable direct model comparison. In the integrated atlas, we identified eight integrated fibroblast cell states (IFS) by performing unsupervised clustering. Each study contributed to all IFS (Supp. Fig. 5A). To functionally characterize the IFS, we derived state markers via differential expression analysis (Fig. 2D, Supp. Fig. 5B, Supp. Table 1).

**Figure 5.**
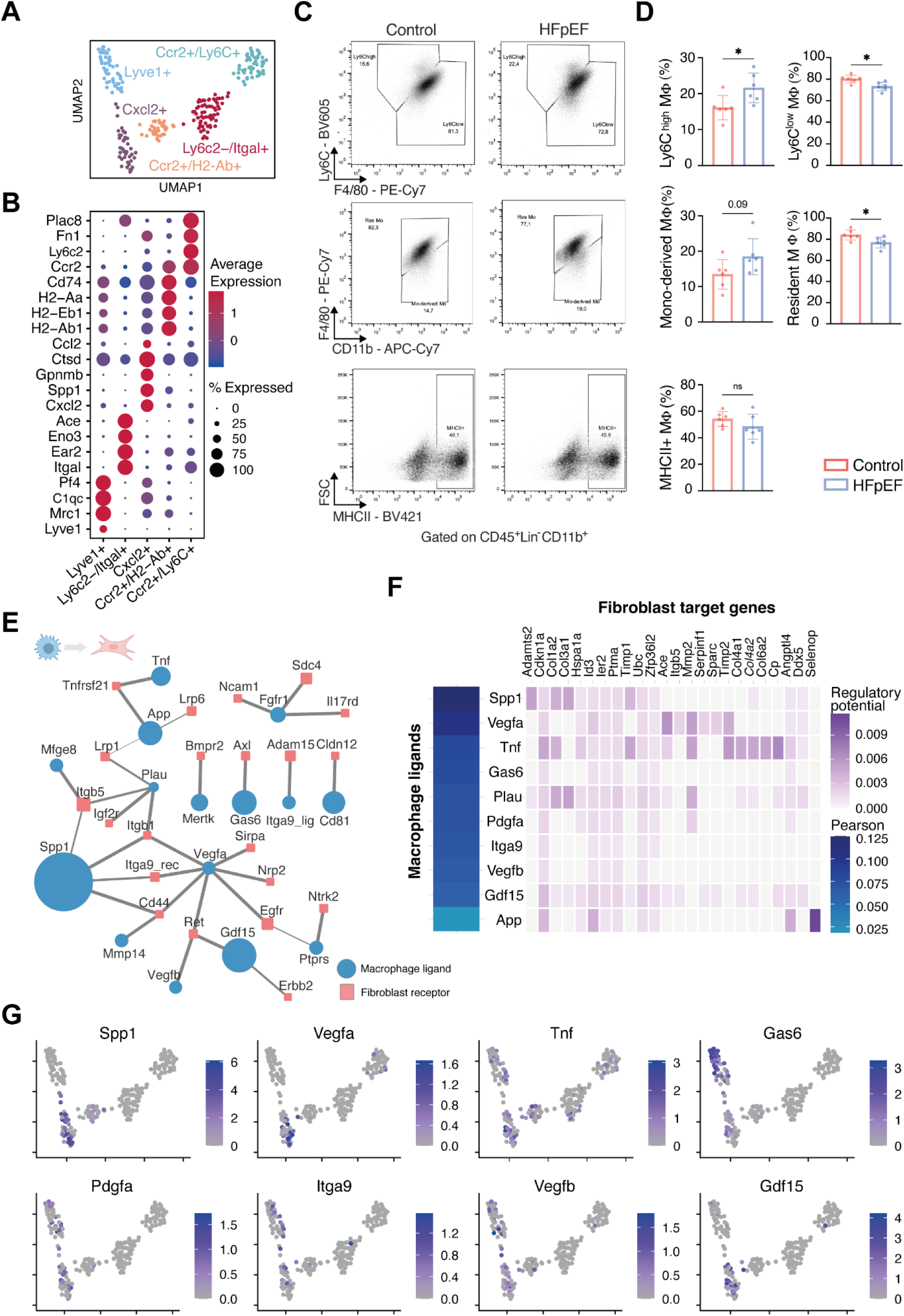
Macrophage engagement in HFpEF. A) UMAP embedding of reintegrated and reclustered macrophages from control and HFpEF mice. B) Top marker gene expression for macrophage and monocyte subtypes. C) Representative flow cytometry plots of Ly6C^high/low^ monocytes and macrophages (MՓ) (upper row), monocyte-derived/ resident MՓ (center row) and MHCII+ MՓ in HFpEF vs control mice. Cells were gated on CD45^+^Lin^+^CD11b^+^ cardiac cells. D) Quantification of flow cytometry results. Statistical analysis using t-test, bar graphs indicate mean±SD, n= 6/group *p<0.05, ns= not significant. E) Ligand-Receptor network based on LIANA, receptors in fibroblasts shown in red, blue depicts ligands from macrophages. Node size visualizes effect size of upregulation in HFpEF mice, edge width visualizes HFpEF specificity (see methods). F) Pearson correlation of top predicted ligands in HFpEF (from E) in NicheNet (left panel). Top NicheNet ligands and their regulatory potential with fibroblast target genes (right panel). G) Top ligands and their normalized expression visualized in UMAP embedding in macrophages.

First, we aimed to identify which of these IFS constitute cardiac specific fibroblasts. For this, we compared IFS markers with markers of fibroblast states identified in a cross organ fibroblast atlas (Supp. Fig. 5C)^19^. IFS 0 (Col15a1+), 3 (Comp+) and 4 (Pi16+) displayed high marker overlap (hypergeometric test p<0.01) which suggested that these states might represent fibroblast phenotypes shared across organs. Conversely IFS 1, 2, 5, 6 and 7 displayed weaker or ambiguous associations and could represent rather cardiac specific fibroblast phenotypes.

We expected that IFS could represent functional niches (i.e. specialization of fibroblasts to fulfill certain tissue functions). We characterized functional niches (Fig. 2E) by performing pathway activity (Fig. 2F) and gene set enrichment analysis (Fig 2G, Supp. Fig. 5F). **IFS 0** fibroblasts were the most abundant cell type in every dataset (Supp. Fig. 5D) and have been described as homeostatic fibroblasts that are characterized by Col15a1 and Dpep1 expression^19^. **IFS 4** fibroblasts were characterized by Pi16 expression and constitute adventitial stromal cells that might accomplish a reservoir function for downstream fibroblast differentiation^19, 21^. The **IFS 3** can be termed matrifibrocytes and are characterized by Cilp, Thbs4, Comp and Postn expression^13, 20^. Pathway analysis indicated that IFS 3 demonstrated highest TGFβ activity (Fig. 2F), which highlighted the pro-fibrotic potential of this IFS. Extracellular matrix (ECM) remodeling is a major operation of fibroblasts and was assessed by enrichment of ECM related gene sets^22^, suggesting that **IFS 0** and **IFS 3** fibroblasts are the main ECM producers: both were characterized by expression of collagens and core matrisome related genes, while IFS 0 uniquely expressed genes associated with the basement membrane (e.g. Col4a1, Lamb1, Hspg2, Col15a1)(Fig. 2G). We identified three IFS with inflammatory profiles: **IFS 2, IFS 6** and **IFS 7**. **IFS 2** appeared to be a heterogenous group of fibroblasts that are partly characterized by Acta2 and Actb expression which constitute myofibroblast characteristics, as well as pro-inflammatory genes involved in antigen processing and representation (Psmd8, Psma6, Vamp8) and Chaperonin containing T-complex polypeptide (CCT) genes (CCT3, CCT7, CCT4, CCT8) that have been associated with proliferative and fibrotic tissue remodeling^23–25^. Furthermore IFS 2 exhibited highest PI3K pathway activity which has been shown to enable fibroblast migration^26, 27^. **IFS 6** fibroblasts were characterized by pro-inflammatory NFκB and TNFα signaling (Fig. 2F) and cytokine expression of Ccl2, Cxcl5 suggested that IFS 6 participates in immune cell attraction. **IFS 7** cells formed a small cluster that every study contributed to with a comparatively small number of cells (Supp. Fig. 5D) and was characterized by JAK-STAT activity and interferon-γ related gene expressions (Ifit3, Isg15). The JAK-STAT pathway has been linked to fibroblast activity in rheumatoid arthritis^28, 29^ and osteoporosis^30^ but its function in cardiac fibroblasts remains unclear. **IFS 5** was characterized by, among others, Wif1 and Dkk3 expression. In the heart, Wif1+ cells were previously shown to localize at the cardiac valves and their adjacent hinge regions and are suggested to be specialized on this spatial niche^31^. **IFS 1** was characterized by expression of typically secreted gene products including insulin-like growth factor 1 (Igf1) and fibrinogen-like protein 2 (Fgl2), which can control cardiomyocyte growth^32, 33^, next to the Igf-function regulators insulin-like growth factor-binding proteins (Igfbp3, Igfbp4)^34^, glycoproteins like fibulin-1 (Fbln1), extracellular matrix protein 1 (Ecm1) and matrix-gla protein (Mgp).

In summary, the phenotype atlas of murine cardiac fibroblasts can be broadly categorized as a set of eight fibroblast states, characterized by distinct key molecular programmes including ECM remodeling (IFS 0, 3), immune modulation (IFS 2,6,7), secretion (IFS 1), and presumably tissue homeostasis (IFS 1, 5).

### Distinct fibroblast signature of HFpEF

To functionally compare fibroblast activation between study models, we performed differential gene expression analysis between control and disease fibroblasts for each study independently to avoid cross study batch comparison. Since the MI study included multiple timepoints, we separated the samples by calculating signatures of early (days 1, 3, and 5) and late (days 7 and 14) remodeling. The resulting signatures contained a small set of upregulated (Timp1, Col1a1, Col1a2, Loxl1 and Sparc) and downregulated genes common to all disease models (Fig. 3A, Supp. Table 2). Between the HF models we found little overlap regarding the respective differentially expressed genes, except for AngII and late MI signatures (Fig. 3B). Since only a few genes were shared between disease signatures, we asked whether the direction of gene expression regulation, determining whether a gene’s activity is increased (upregulated) or decreased (downregulated), is nevertheless consistent between HF models. We correlated fold change regulation of disease signatures between studies and found the strongest correlation between AngII and late MI fibroblasts.

Interestingly, the HFpEF signature did not correlate with AngII while displaying weak agreements with early and late MI, indicating disease-specific fibroblast activation patterns.

To further elucidate the differences of these fibrotic features, we characterized signatures by enriching annotated gene sets from the MSIG database. Fibrosis signatures across models contained major ECM related gene sets (Fig. 3D), indicative of a common profibrotic task. The HFpEF signature was uniquely characterized by heat shock factors, protein glycosylations, basement membrane and laminin components, but contained less components related to elastic fibers than AngII and MI (Fig. 3E). Next, we used fold change regulation of regulon genes to infer upstream transcription factor (TF) activities (Fig. 3F). Among others, Hsf1, Ppar-ɑ and Ppar-ɣ are suggested to be relevant TFs specifically in HFpEF fibroblasts and could constitute important mediators of metabolic stress response. In addition, Hif1ɑ activity was found in HFpEF and, as expected, in early MI fibroblasts. In HFpEF, hypoxia may occur in obesity related tissue stress^35^, but its impact on cardiac fibroblast function in HFpEF is unknown. All models displayed high Smad3 activity, which is in line with the common knowledge that this TF is an important driver of cardiac fibrosis, possibly via TGFβ signaling^36^. Indeed, when comparing pathway activities (Fig. 4G), TGFβ was active in all HF models, however, strongest activity was found in AngII and late MI models while in early MI fibroblasts proinflammatory TNFα, NFκB, as well as hypoxia, and JAK-STAT pathways were induced. In HFpEF, besides TGFβ, the hypoxia^37, 38^, TNFα^39^ and p53^40^ pathways were predicted to be activated in fibroblasts.

Besides the upregulation of main ECM components in HFpEF (e.g. Col1a1, Col1a2, Col4a1, Sparc, Pcolce), we found collagen cross linking enzymes (Loxl1, Loxl2), metabolic and inflammation related genes to be induced (e.g. Angptl4, Ace, Dpep1, If205 and Ccd80) (Fig. 3H). Col4a1 is an important component of the basement membrane and its accumulation over time in the HFpEF model was confirmed by immunofluorescence stainings and indicated a collagen IV pattern of interstitial sheathing of cardiac cells (Fig. 3I). Angiopoietin-like 4 (Angptl4) is a lipoprotein lipase inhibitor that was barely expressed in control fibroblasts, but strongly induced in HFpEF. It is known to be regulated via Ppar-ɑ and Ppar-ɣ in other contexts^41^. Hence, Angptl4 could constitute an important indicator of metabolic stress in fibroblasts in HFpEF. Qualitative protein staining of Angptl4 confirmed upregulation especially in the cardiac interstitium (Fig. 3J).

While the common gene expression patterns between HF models related to TGFβ and Smad3 activity together with upregulation of ECM genes, distinctive HFpEF fibroblast activation patterns included upregulation of Angptl4 and other markers of metabolic stress, basement membrane genes, and activation of proinflammatory pathways and TFs.

### Compositional and transcriptional shifts in cardiac fibroblasts

In the previous sections, we characterized integrated fibroblast states (IFS) together with their possible functional niches and interpreted model-specific disease signatures. To combine both perspectives, we investigated how fibroblasts from different IFS contributed to the model specific cardiac remodeling. This could help us understand the division of labor between fibroblast states and compare study models from a cell population perspective.

We conceptualized different patterns of gene expression with respect to cell states that lead to an upregulation of a disease signature (Fig. 4A). First, we distinguished between compositional and transcriptional shifts^42^. The former describes a relatively stable expression within a cell state where the disease signature upregulation is caused by an increase in the proportion of that state. On the other hand, a transcriptional shift constitutes an upregulation without a compositional increase. Here, we propose to differentiate between upregulation focused within a state (state-dependent) and within many or all states (state-independent). We will use these terms to broadly describe gene expression patterns from different HF models, however, we acknowledge that these categories are not exclusive.

To investigate compositional shifts, we calculated compositional changes of IFS between control and diseased mice per study (Fig. 4B). In HFpEF, composition changes were very small and only IFS 0 and 6 expanded slightly (label permutation p-value <0.05), while in early MI the highest compositional dynamics were observed with expansion of IFS 2, 3, 5 and 6. Late MI remodeling displayed similar characteristics as the AngII model with an increase of IFS 3 and 2. To support that these compositional shifts were associated with the disease signatures we assessed the overlap of genes between IFS markers and disease signatures (Fig. 4C). We found that IFS 0 shared markers with the HFpEF signature while IFS 3 with AngII, late and early MI signatures; IFS 2 and 6 with early MI signature only (hypergeometric test, p<0.05). This suggested that a compositional shift was happening in all mouse models, but with different emphases of IFS across the HF models. No other model shared the importance of IFS 0 with HFpEF, which could possibly be a unique feature of HFpEF fibrosis.

To investigate transcriptional shifts, we quantified how well fibroblasts within the same IFS can be distinguished regarding their control and disease label as a metric for a transcriptional shift (Fig. 4D, Supp. Fig. 6A,B) (see methods). In general, the signatures were increasingly expressed across IFS and thus a transcriptional shift was apparent in every study model (Area under the receiver operator curves, AUROCs > 0.5). The highest differences were achieved in the MI (early and late) models compared to HFpEF and AngII models which could indicate that the latter displayed a less pronounced transcriptional shift. This might be explained by acute tissue injury after MI as opposed to the chronic stimuli of AngII administration or HFpEF diet. In addition, a different hierarchy of IFS responsiveness was observed, indicating that the transcriptional shift is partially state dependent: While the highest transcriptional shifts in HFpEF were found in IFS 7 and 0, the other models consistently displayed IFS 3 and 7 as the most responsive states (Figure 4D). Furthermore, IFS 5 fibroblasts were poorly responsive in all study models and thus probably less relevant for disease remodeling which might relate to their reported localization particularly at the cardiac valves.

**Figure 6.**
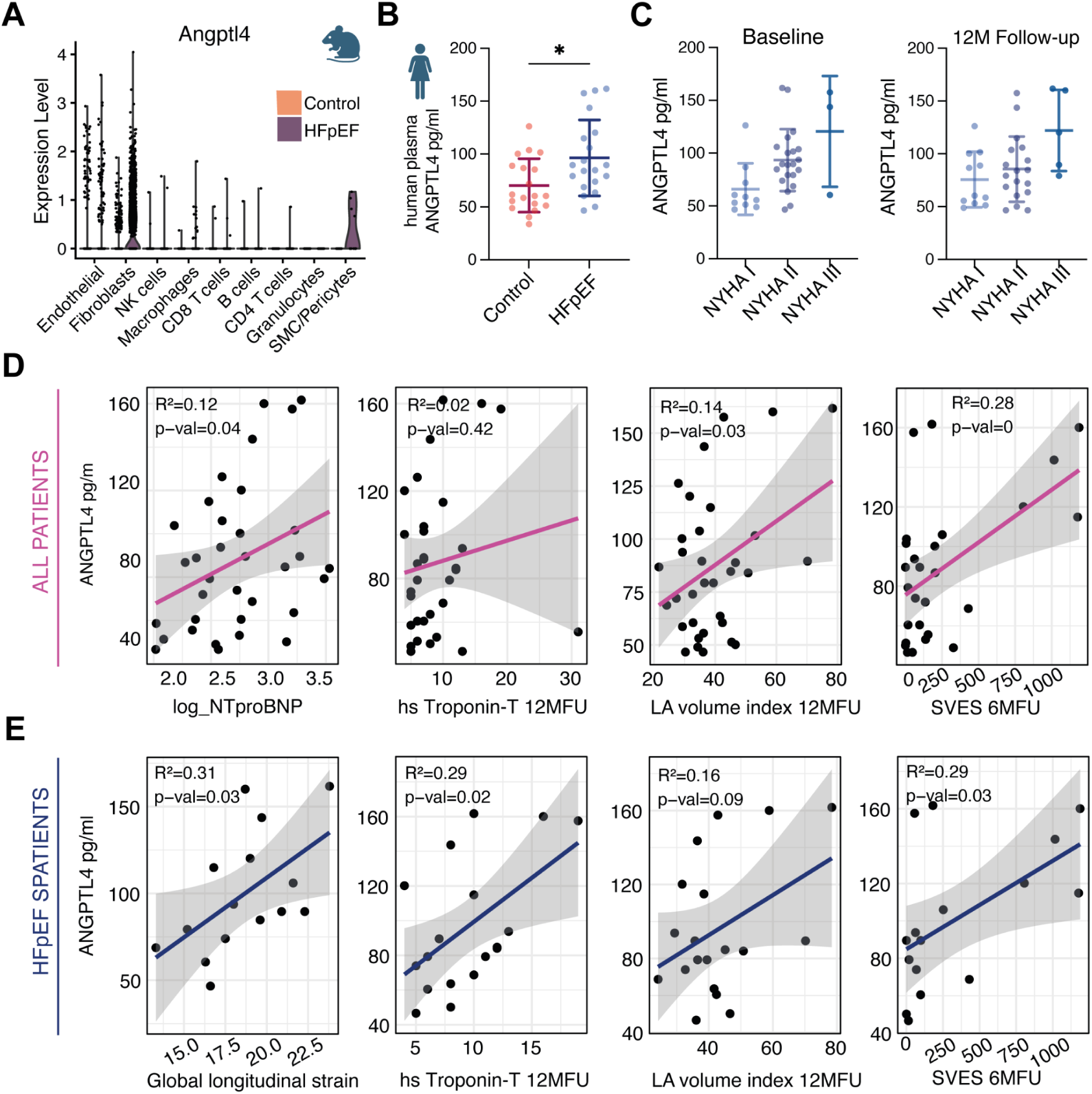
ANGPTL4 expression is increased in murine HFpEF hearts and in plasma of HFpEF patients. A) Angptl4 normalized gene expression among different cardiac interstitial cell types derived from the scRNAseq data showing control (left columns) and HFpEF (right columns) separated per cell type. B) Circulating levels of ANGPTL4 in human plasma samples of HFpEF and age matched controls measured by sandwich ELISA. n=19/20, Mann-Whitney U test, *p<0.05. C) ANGPTL4 plasma levels in relation to NYHA functional class of all recruited patients. ANOVA, p-value <0.05, n= 10/21/3 in baseline and n= 11/18/5 in 12 months (12M) follow-up. D) Correlation of clinical parameters to ANGPTL4 circulating levels in all patients (control and HFpEF) and E) as subanalysis only in HFpEF patients using simple linear regression. p-val indicates p-value. hs= high sensitivity, LA= left atrial, MFU= months follow-up, SVES= supraventricular extrasystoles. Plots in B,C display mean±SD.

In summary, we found that fibroblasts from IFS that did not display compositional shifts, nevertheless, contributed to the remodeling by upregulating respective disease signatures. However, fibroblasts from IFS that do display compositional shifts displayed also a high transcriptional shift, suggesting that both concepts are biologically closely related. We concluded that IFS that displayed i) state marker overlap, ii) compositional increase and iii) a high within-state-transcriptional-shift, represented the desired functional niches and thereby the prioritized states in each HF model. Those states were IFS 0 in HFpEF, IFS 2, 3 and 6 in early MI and IFS 3 in late MI and AngII. However, besides this prioritization, all states apart from IFS 5 partook in cardiac remodeling.

### Decomposing disease signatures and state dependency

After characterizing transcriptional and compositional shifts in the HF models, we next aimed to decompose disease signatures regarding their state dependency. To quantify this dependency we calculated eta² values (see methods). In HFpEF fibroblasts, genes related to the basement membrane (Lamc1, Lamb1, Col4a1, Nid1) yielded highest eta² values suggesting a state dependent expression. Metabolism (Angptl4, Ech1, Man2a, Acaa2) and fibrosis associated genes (Col1a1, Col1a2, Timp1, Mmp1) displayed rather state independent expression patterns (Fig. 4E). This indicated that basement membrane remodeling might be a functional specialization of fibroblasts, while the upregulation of metabolic and protein stress together with non basement membrane ECM markers are a more general gene program of fibroblasts in HFpEF.

To compare these expression patterns between HF models, we calculated eta² values for state (indicative of state dependency) and group labels (indicative of an upregulation) for all HF models separately (see methods).

First, we quantified the ratio of both eta² values to compare the general state dependency of disease signatures between HF models (Fig. 4G). The two chronic models (HFpEF & AngII) displayed a more state-dependent transcriptional shift compared to the MI (late & early) fibroblasts (Wilcoxon test, p < e10), suggesting that the state dependent fibroblast response might be a characteristic of chronic remodeling.

Second, we compared the state dependency of single genes between HF models (Fig. 4H). State markers like Dkk3 (IFS 5) or Pi16 (IFS 4) displayed high state dependency and low group dependency in all HF models, serving as examples for genes that are state markers but without disease involvement. Angptl4 was exposed as a state independent marker, specific for HFpEF. Postn displayed high group and state related variance in all HF models except HFpEF, and thus represented a crucial marker with high disease association and state dependency. Next, we focused on the core intersection of upregulated genes in all HF models and assessed whether their regulation regarding state dependency might differ between HF models (Fig 4H). We found that most genes were regulated state dependently. Interestingly, Col1a1 and Col1a2 were expressed state-dependently in all HF models, except HFpEF (Fig. 4G). Collagen I is a main ECM component and crucial for integrity and stiffness of fibrotic tissue and it has been reported that matrifibrocytes in the heart are responsible for the deposition of collagen I^43^. Our findings could indicate that fibrosis due to collagen I deposition in the early HFpEF model might not be related to a state dependent task since matrifibrocytes are not activated yet and the early fibrosis is achieved by a collagen production by fibroblasts of all phenotypes.

To demonstrate how the transcriptional shifts of the discussed key genes relate to IFS, we quantified within state regulation of single genes (AUROCs) (Supp. Fig. 6C) and found that Col1a1 and Col1a2 were upregulated in almost all IFS across models, showing highest upregulation within IFS 3 in non HFpEF models. Col4a1 and Col4a2, although state dependent expressed in all models, displayed a high transcriptional shift in most IFS in HFpEF. This further elucidates that genes that are state-dependently expressed between fibroblasts (such as collagen IV in IFS 0 or collagen I in IFS 3) are also upregulated by other IFS but only in the respective disease context. In addition, Angptl4 displayed low state dependent variance and a high transcriptional shift in all IFS in HFpEF, possibly rendering it a key marker of state independent metabolic fibroblast stress.

Differential gene expression analysis can be confounded by background gene expression in single-cell transcriptomics that could be associated with increased cell dissociation in diseasesed tissue affecting contrast comparison. Other cell types in our single-cell data could not be separated on the basis of the discussed disease signatures, suggesting that the discussed transcriptional shifts were not confounded by background expression (Supp. Fig. 6D). Furthermore, the low correlation between HFpEF and other HF signatures (from Fig. 3B,C) caused other disease signatures to fail to separate HFpEF fibroblasts from control.

### Corroborating fibroblast signatures in humans and mice

In the previous section we established the IFS prioritization by the different HF models. To explore whether this IFS to HF phenotype association could be recovered in humans, we curated myocardial bulk transcriptomic signatures acquired from HFrEF and HFpEF patients. For HFrEF, we relied on a meta-analysis of a total of 653 patients with end stage heart failure^44^ For HFpEF, due to limited by data availability, we re-analyzed data from 5 patients that underwent coronary artery bypass graft surgery and met the echocardiographic and diagnostic criteria for HFpEF^45^. We selected top upregulated genes from both bulk resources and performed overrepresentation analysis with the fibroblast disease signatures (Fig. 4I, left panel). The murine AngII and late MI signatures displayed a significant overlap with the human HFrEF bulk reference, while murine HFpEF signatures were enriched in the human HFpEF bulk reference (hypergeometric test, p<0.05). Next, we addressed whether this intersection of disease signals between mouse and human could also be recovered for IFS markers (Fig. 4I, right panel). We found that markers for the IFS 3 (matrifibrocytes) were overrepresented in the human HFrEF signature, in agreement with recent reports from human single-cell studies^46, 47^. In the human HFpEF signature, only IFS 0 state markers were overrepresented. This could possibly suggest a relevance of IFS 0 and its functional niche for human HFpEF. In general, the presented fibroblast signatures from AngII, MI and HFpEF, as well as the IFS prioritizations of models are partially conserved across species (mouse to human) as well as across data modalities (single-cell to bulk RNAseq).

Matrifibrocyte activation is a crucial event in cardiac fibrosis and we further explored the role of this event in the murine HFpEF model. First, we compared the protein expression of myofibroblast and matrifibrocyte markers, such as Cilp, in HFpEF with a MI model (Supp. Fig. 7A). The matrifibrocyte marker Cilp (Cartilage Intermediate Layer Protein) is an ECM protein abundant in articular cartilage and has been implicated in cardiac fibrosis before^48, 49^. Immunohistological staining of Cilp displayed moderate perivascular protein expression in myocardial tissue of 10 weeks HFpEF compared to control mice, while strong expression was observed after MI (Supp. Fig. 7B). Fibroblast activation protein (Fap) was introduced as a marker for myofibroblast activation in cardiac fibrosis^50, 51^. Fap stainings demonstrated that its expression could only be observed in the MI fibrotic zone (Supp. Fig. 7B). We further assessed Fap expression within the whole murine organism after 15 weeks of HFpEF diet by PET-CT based ^68^Ga-FAPi-46 uptake (Supp. Fig. 7C), yielding no relevant expression patterns across organs. This data indicated that based on myofibroblast and matrifibrocyte markers (Fap and Cilp, respectively) the HFpEF model is not associated with a strong response of these cell states.

Second, we explored myocardial gene expression after 10 and 15 weeks of HFpEF diet (compared to seven weeks in the single-cell data only containing interstitial cells) (Supp. Fig. 8A). We contrasted both timepoints to control mice and found that the HFpEF, MI and AngII disease signatures all enriched at 10 and 15 weeks (Supp. Fig. 8C). When comparing IFS markers, we found that IFS 0, 1, 3, 5 and 7 enriched significantly (linear regression, p<0.05) with IFS 3 yielding the highest enrichment score (Supp. Fig. 8D). This could indicate that a matrifibrocyte activation as reflected by IFS 3 marker upregulation might occur later than seven weeks in the HFpEF model. Nevertheless, IFS 0 and HFpEF disease signature upregulation could be recovered as well, suggesting that the described cellular pathways are partially coincidental events in the murine model over time.

### Macrophage activation in single-cell transcriptomics and flow cytometry

The cellular and molecular pathways that lead to the activation of fibroblasts in HFpEF are unknown. One possible role could be attributed to macrophages which have been discussed as a crucial modulator of fibroblast activity in HFpEF^12, 52^. Our single-cell data suggest evidence for macrophage involvement (Fig. 1F), therefore we further investigated macrophage phenotypes in the HFpEF model.

We identified four cardiac macrophage and one Ccr2+/Ly6c+ monocyte cluster (Fig. 5A), the latter expressed high levels of marker genes of inflammatory monocytes (Ly6c1 and Ccr2) and fibrosis mediating genes (Fn1, Thbs1 and Vim). The macrophage cluster differed in marker expression of monocyte-derived or resident macrophage related genes (Fig. 5B). To estimate reliable macrophage compositions without a relevant sampling-error, which might limit interpretation of our single-cell data due to low cell counts, we performed flow cytometry experiments of the entire ventricular tissue. The flow cytometry data reveal a significantly expanding proportion of pro-inflammatory Ly6C^high^ monocytes/macrophages next to decreased Ly6C^low^ macrophages. Gating macrophages according to their F4/80 and CD11b expression showed a significantly reduced proportion of resident (not monocyte-derived) macrophages, potentially driven by expanded monocyte-derived macrophages, which did not reach statistical significance (Fig. 5C-D, Supp. Fig. 9A). Total counts of tissue leukocytes, granulocytes and macrophages subsets did not differ significantly (Supp. Fig. 9B-E). To analyze whether the shift to pro-inflammatory macrophages is related to a splenic activation as a major source for myeloid cells in acute tissue injury^53, 54^, we performed flow cytometry analysis of spleen and also peritoneal macrophages as potentially contributing inflammatory compartments following HFpEF diet. Neither splenic nor peritoneal macrophages showed a significant induction of pro-inflammatory subsets, such as Ly6C^high^ spleen or small peritoneal macrophages^55^ (Supp. Fig. 9F-J). Taken together, next to expanding Cxcl2+, Ccr2+/H2-Ab+ and Lyve1+ macrophages in single-cell transcriptomics (Supp. Fig. 9K), we observed local changes in HFpEF cardiac tissue towards a pro-inflammatory monocyte/ macrophage composition, but not systemically in splenic and peritoneal compartments.

### Cell-cell communication between Macrophages and Fibroblasts

The observed fibroblast activation and inflammatory response of macrophages lead us to hypothesize about potential cellular communication between both cell types. We used LIANA^56^ to score ligand-receptor (LR) interactions between macrophages and fibroblasts in control and HFpEF mice (see methods, Supp. Fig. 10A). Top predicted LR pairs were upregulated in HFpEF (Supp. Fig. 10B) and included Spp1 binding CD44 or Itgb1, and Tnf binding Tnfsrsf21 (Fig. 5E). To identify possible links to the HFpEF fibroblast disease signature we used NicheNet^57^ to assess the regulatory potential of predicted ligands (Fig. 5F, left panel). Spp1 is predicted with regulatory potential affecting core fibrotic genes such as Col1a2, Col3a1, Adamts2 and Timp1, while Tnf ligand might be associated with basement membrane component Col4a1 and Angptl4 regulation (Fig. 5F, right panel). Expression patterns of ligands (Fig. 5G) suggest that Cxcl2++ macrophages could communicate via Spp1, Vegfa, Gdf15, and Plau ligands while Lyve1+ macrophages might secrete Gas6 and Pdgfa ligands. Tnf and Vefgb ligands are expressed in both states. We assessed regulatory patterns of these predicted LR pairs in the other HF models and found that Spp1 was induced in HFpEF and early MI and thus constitute a mediator of the inflammatory axis that is not involved in matrifibrocyte activation (Supp. Fig. 10D, E).

### Circulating ANGPTL4 levels in HFpEF vs. non-HFpEF patients

Identifying traceable markers of fibroblast activation in humans could help to assess and possibly target HFpEF remodeling at an early stage. We described the expression pattern of Angptl4 as a state independent marker of fibroblast activation in murine HFpEF with little expression in other interstitial cell types (Fig. 6A). We further confirmed A upregulation on protein level and reported Ppar-ɑ and Ppar-ɣ to be active TFs as possible upstream regulators, together with a regulatory potential via macrophage based TNFɑ. Angptl4 is functionally linked to inflammation, metabolism and fibrosis^58^. Thus, we hypothesized that ANGPTL4 might be a promising candidate to be involved in HFpEF pathophysiology and evaluated whether ANGPTL4 could serve as a biomarker detectable in human plasma.

We analyzed circulating levels of ANGPTL4 in 20 plasma samples of HFpEF and 20 non-HFpEF (control) patients. All patients were diagnosed for symptomatic atrial fibrillation and screened for HFpEF by echocardiography, stress echocardiography, NT-proBNP, and HFA-PEFF-score^59^. Plasma samples were analyzed by ELISA, which revealed significantly higher circulating ANGPTL4 levels in HFpEF (Fig. 6B). ANGPTL4 levels increased significantly in higher NYHA stages in all patients (Fig. 6C) and correlated significantly with NT-proBNP, but not with high-sensitivity troponin T (Fig. 6D). In a subanalysis of the HFpEF cohort, high ANGPTL4 levels related positively to counts of supraventricular extrasystoles in holter ECGs and left atrial volume index (biplane, ml/m^2^), at 6-and 12-months follow-up, respectively (Fig. 6D-E), but not at baseline indicating an association with disease progression as it is known that left atrial dilatation and supraventricular arrhythmias are associated with HFpEF severity^59^. Exclusively in HFpEF, but not in control patients, plasma ANGPTL4 was associated with troponin T levels and global longitudinal strain (Fig. 6E). This data suggested that plasma levels of ANGPTL4 might be mechanistically linked to disease characteristics of human HFpEF and its therapeutic or prognostic potential should be further evaluated.

## Discussion

In this study, we provided a first comprehensive characterization of interstitial cardiac remodeling in a two-hit HFpEF mouse model on single-cell level. Deterioration of cardiac diastolic function was accompanied by increased perivascular fibrosis in murine HFpEF hearts. This phenotype was associated with a pro-fibrotic gene program in fibroblasts. By integrating single-cell atlases of two additional murine HFrEF models, we identified conserved fibroblast states across models and derived common and unique functional characteristics of fibroblast activation in HFpEF compared to HFrEF. We corroborated disease signatures in human transcriptome data, and suggested possible involvement of macrophages in the activation of fibroblasts in HFpEF. Finally, we suggested that Angptl4, which was elevated in HFpEF patients and related to disease characteristics and functional capacity, could serve as a potential biomarker for disease severity in patients.

Fibroblast activation and cardiac fibrosis are hallmark features of ventricular remodeling in heart failure (HF). However, disease specific molecular patterns of fibroblast function in different types of HF are unknown. Here, we compared fibroblast activation in early HFpEF (two-hit model), renin-angiotensin-aldosterone system activation-induced HFrEF (AngII model) and early and later ischemic HFrEF (myocardial infarction model) to identify common fibroblast phenotypes across models. Among fibroblasts, different phenotypes are assumed to represent functional and/or spatial niches^60, 61^. However, a consensus and nomenclature of cell states has not been accomplished yet, in part due to shortcomings of the concept of cell states attempting to i) categorize a continuity and ii) distinguish between a more transient functional nature of a state or a cell differentiation^60^. By integrating multiple studies, we provided a catalog of conserved cardiac fibroblast cell phenotypes in heart failure, possibly representing the hallmarks of cardiac fibroblast function, including ECM production (IFS 0, 3), secretory function (IFS 1), immune system modulation (IFS 6,2,7), migration (IFS 2) and tissue homeostasis (IFS 4, IFS 5).

Despite this functional diversity, ECM remodeling and collagen deposition was a common fibroblast task across models, reflecting fibrosis as disease characteristic in each HF model. In HFpEF fibroblasts, metabolic stress, heat shock proteins and glycosylation of proteins were accompanied by upregulation of ECM components, in particular basement membrane compounds. The basement membrane represents a highly active ECM that underlies many cell types such as ECs and SMCs and provides a scaffold that connects cardiomyocytes to the ECM^62^. Functionally, it plays an important role in angiogenesis, mechanotransduction and cell differentiation^63^. The role of the basement membrane in HFpEF has not been sufficiently explored yet, but its modulation of laminins has been suggested to cause gene expression changes in cardiomyocytes related to increased stiffening^64^. Interestingly, HFpEF shared proinflammatory features like hypoxia and TNFα pathway activity with early MI fibroblasts, while late MI and AngII fibroblast displayed highest TGF□ activity. In addition, we observed an expansion of pro-inflammatory Ly6C^high^ monocytes/macrophages in HFpEF hearts and predicted a mutual activation occurs in the cross-talk with fibroblast via Spp1 and TNFα. A potential therapeutic benefit of TNFα inhibitors in HFpEF has not been investigated and deserves further investigation.

Single-cell transcriptomics enable a deeper characterization of cell type function in disease by regarding the division of labor between cells and their functional or spatial niches^65, 66^. We demonstrated that fibroblast activation is a mixture of compositional and transcriptional shifts in all HF models with the strongest transcriptional shift following MI, indicating that acute tissue injury induces more population wide cell responses. Besides these programs, prioritized states were identified in each model, suggested by high transcriptional shifts co-occurring with compositional shifts in these states. We found that during early MI, migratory myofibroblasts (IFS 2), matrifibrocytes (IFS 3) together with other proinflammatory states (IFS 7 and 6) were prioritized, in contrast to AngII and late MI which mainly exerted fibrosis via matrifibrocytes. The early HFpEF-associated fibrosis might differ from these respective HFrEF-like remodeling processes, as we found little disease signals of matrifibrocytes, but identified homeostatic IFS 0 fibroblasts and basement membrane remodeling as key characteristics. Collagen I deposition is crucial for pro-fibrotic ECM remodeling and has been described as a characteristic of matrifibrocyte activity. In early HFpEF, the division of labor of collagen I synthesis was shifted from a state dependent task of matrifibrocytes to a general fibroblast task. This could possibly be associated with the extent of the observed cardiac fibrosis. A subsidiary role of matrifibrocyte activation in HFpEF has been suggested before by the evidence of ECM production in the absence of TGFβ signaling and presence of metabolic stimulation^10^. Our data further supported this hypothesis by demonstrating that no relevant FAP expression was observed in HFpEF hearts in contrast to the previously described upregulation in acute MI and AngII/PE^50, 67^. The translational potential of our findings to human disease is highlighted by the corroboration of the described fibrotic signatures in myocardial transcriptomes of human HFrEF and HFpEF. Therefore, it can be assumed that the established treatment options for HFrEF that improve maladaptive cardiac fibrosis, such as inhibitors of the renin-angiotensin-aldosterone system^68, 69^, failed to obtain this therapeutic potential for HFpEF possibly due to distinctive fibroblast activation patterns different from HFrEF with less implications of matrifibrocyte and myofibroblast activation and higher impact of metabolic alterations.

Regarding metabolic changes in HFpEF interstitial cells, we observed a strong induction of Angptl4 as opposed to very decent RNA and protein levels in control hearts. Angptl4 is linking metabolic, inflammatory, and fibrotic mechanisms by acting as a secreted matricellular protein and by controlling metabolism by inhibiting intracellular lipoprotein lipase in divergent tissues^58^. Induction of Angptl4 gene expression via Ppar-β/δ in cardiomyocytes, triggered by dietary fatty acid administration, exhibited a protective effect against lipid overload and subsequent fatty acid–induced oxidative stress^70^. Next to its important role as metabolic regulator in cardiomyocytes, Angptl4 is expressed in mesenchymal stem cells and might balance innate immune responses such as promoting anti-inflammatory macrophage polarization following MI^71^. In non-cardiac endothelial cells hypoxia enhanced Angptl4 expression and full-length Angptl4 deposition was observed in the subendothelial perivascular space of ischaemic hindlimbs and suggested to be originated from endothelial cells, even though the authors did not consider other potential sources, like fibroblasts^72, 73^. Here, we suspected fibroblasts as the central interstitial cell type for Angptl4 secretion into the ECM. However, the functional role of Angptl4 expression in cardiac fibroblasts remains barely investigated. Our single-cell fibroblast data predicted an activation of transcription factors Ppar-ɑ, Ppar-ɣ, Hif1ɑ and hypoxia pathways in murine HFpEF. In our small patient collective with atrial fibrillation circulating ANGPTL4 was related to reduced functional capacity, NT-proBNP and HFpEF disease progression by relating to left atrial dilation and burden of supraventricular extrasystoles at follow-ups, but correlated positively with global-longitudinal strain. In contrast, murine studies showed that recombinant Angptl4 attenuated Ang II-induced atrial fibrillation and atrial fibrosis via Sirt3, Ppar-ɑ, and Ppar-ɣ, signaling pathways^71, 74^. Next to conflicting murine reports about adverse effects, such as reduced cardiac function in Angptl4 overexpressing mice, an association of high circulating Angptl4 levels with the risk of coronary artery disease, atherosclerosis and type 2 diabetes in humans was described^71, 74^. While Angptl4 has been shown to play a major role in regulating intracellular metabolic adaptation, its effects appear to be diverse and context-dependent, with both beneficial and partially detrimental functions across different cell types and compartments. Further research is needed to fully elucidate its mechanistic role in cardiac fibroblasts and its impact on HFpEF disease traits and outcome, with the aim of identifying novel therapeutic approaches based on the distinct pathomechanisms.

The main limitations of our study relate to the sample size of the single-cell experiment. Subtle disease changes, such as gene programs occurring in more rare cell types or cell states, were probably not detectable. At the same time, our statistical approach for differential expression and composition analysis might result in a higher rate of false positives than more robust approaches that rely on higher sample size. However, multiple confirmation experiments suggested that disease signatures were reproducible in other data. Our study design focused on early changes of the murine HFpEF model. As a longer dietary regimen might lead to further disease progression, we cannot provide insights into potential dynamics of the reported cellular disease signatures on single-cell level. A potential role of matrifibrocytes during later stages of the HFpEF model was suggested by upregulation of matrifibrocyte markers in bulk transcriptomics. However, it is unclear which time point of the murine model represents most closely human HFpEF. Corroboration in human bulk transcriptome demonstrated that matrifibrocyte markers are upregulated in human HFrEF, but not in HFpEF patients. Additional validation of these findings in larger human HFpEF studies could not be accomplished, due to the small number of publicly available datasets of gene and protein expression in human HFpEF.

Common fibrotic pathways are active across pathologies, organs and species and include hallmark signaling mediated by TGFβ, integrins, cytokines and vasoactive substances^75^, resulting in increased ECM production and reparative tissue replacement. Pharmacomodulation of these major fibrotic axes has been mainly unsuccessful in the past, partially because of their fundamental impact on global tissue homeostasis. As evidence on more granular differences in fibrotic signaling, fibroblast phenotypic heterogeneity and coordinated functionality is accumulating^19, 76^, better targeted antifibrotic therapies might come within reach^77^. To summarize, we provided a first description of adverse interstitial remodeling in HFpEF at a single-cell level. Our work generated new insights into distinct and common features of cardiac fibrosis in heart failure and might serve as a valuable resource for the scientific community to identify disease specific treatment strategies for HFpEF in the future.

## Online Methods

### Animals

All animal experiments were conducted in agreement with the animal welfare guidelines and German national laws. All animal procedures and study protocols were authorized and approved by the responsible authority (permit No. G-252/20 and G-121/21, Regierungspräsidium Karlsruhe, Baden-Württemberg, Germany). C57BL/6N male mice, obtained from Janvier Labs, were used at an age of 10 weeks. Mice were kept at 23 °C ambient temperature and in 12 h light/dark cycle and had unrestricted access to food (D12450B, control diet rodents 5% fat and D12492, rodent 60% high-fat diet for the HFpEF group, Ssniff) and water. HFpEF was induced as reported previously^3^. Briefly, Nω-nitro-l-arginine methyl ester (L-NAME, 0.5 g/l, Sigma-Aldrich), adjusted to pH 7.4, was supplied by the drinking water in light-protected bottles for the indicated time. Fig. 1A (left panel) and Supp. Fig. 1B+D included n=11 controls (7, 10 and 15 week control diet), n= 8 7-week HFpEF, n= 9 10-week and n= 7 15-week HFpEF animals. Acute MI paraffin embedded sections were derived from a C57BL/6N mouse 28 days after minimal-invasive occlusion of the LAD, as described previously^78^. Infarct size of large MI can be gathered from Supp. Fig. 8A.

### Echocardiographic measurements

Transthoracic echocardiography was performed on a VisualSonics Vevo 2100 system equipped with MS400 transducer (Visual Sonics). Left ventricular (LV) parasternal long axis and short axis views at the mid-papillary muscle level were acquired by induction (4 vol%) and short maintenance (0.5–1.5 vol%) of isoflurane anesthesia. LV end-diastolic volume (LVEDV), fractional area change (FAC), LV fractional shortening (FS) and LV ejection fraction (LVEF) were obtained at a heart rate between 500-600 bpm. Parasternal long-axis traces were used to calculate the global longitudinal strain software-and speckle-tracking algorithm-based (VevoStrain software,Visual Sonics). Borders of the endocardium and epicardium were subsequently traced before a semi-automated strain analysis was performed by the software.

For diastolic function, mice were anesthetized under body temperature-controlled conditions and maintenance (1.5-3 vol%) of isoflurane anesthesia aiming to keep the heart rate in the range of 400–450 bpm. Apical four-chamber views were obtained and pulsed-wave and tissue Doppler imaging at the level of the mitral valve performed to record the following parameters: peak Doppler blood inflow velocity across the mitral valve during early (E) and late diastole (A), isovolumic relaxation time (IVRT) and peak tissue Doppler of myocardial relaxation velocity at the mitral valve annulus during early diastole (E’). Analysis was performed with VisualSonics Vevo Lab software, using semi-automated LV tracing measurements for LVEF and FS. All parameters were measured in at least three cycles, and means were presented. GLS measurements could not be performed retrospectively in the animals used for scRNASeq due to bad semi-automated tracing of the images.

### Single-cell RNA sequencing

Sample preparation and sequencing of murine cardiac interstitial cells was performed according to the detailed protocol published previously with only minor modifications^79^. In brief, rapidly after sacrificing the animals by cervical dislocation, the still beating heart was directly placed in ice-cold HBSS where atria and large vessels were dissected. After chopping the heart into small pieces, enzymatic digestion was initiated in two rounds of 15min duration at 37°C using collagenase type II (Worthington Biochemical Corporation, # LS004177). The single-cell suspension was subsequently passed through a 40 µm cell strainer, washed and red blood cell lysis performed. Dead cells were removed by Dead Cell Removal MicroBeads (Miltenyi Biotec, 130-090-101) binding to MACS LS columns (Miltenyi Biotec, 130-042-401). Prior to loading the 10x platform, live nucleated cells (DR^+^, propidium iodine^-^) were sorted using a FACSAria^TM^ IIu (BD) cell sorter. Following washing and resuspending in PBS, cells were counted manually using trypan blue and a Neubauer chamber. We aimed to load about 5,000 cells per lane on a Chromium Next GEM Chip (10x Genomics, 1000127), that was placed into a 10X Chromium Controller (10X Genomics). The cDNA output was amplified and library construction was performed according to the manufacturer’s instructions using the Chromium Next GEM Single Cell 3’ Kit v3.1 (10x Genomics, 1000269) and Dual Index Kit TT Set A (10x Genomics, 1000215). Respective library quantification and quality controls were performed using an Agilent 2100 Bioanalyzer and in addition a Qubit HS Assay. Indexed libraries were equimolarly pooled resulting in two sequencing runs (control1+HFpEF1; control2+HFpEF2) using a High Output kit v2.5 (Illumina, 20024907) and a NextSeq® 550 (Illumina) sequencer.

### Data preprocessing and QC

The resulting single-cell RNA-seq outputs were processed using CellRanger provided by 10x genomics. Count data was processed sample wise with the following filters: >300 Feature numbers, <25% mitochondrial genes, <1% ribosomal genes and >500 RNA counts. Doublet scores were calculated with the R-package *scDblFinder*^80^ and only predicted singlets were kept. We further calculated a dissociation score by estimating expression of dissociation associated gene expression^81^ with Seurat’s^82^ AddModuleScore function and we removed cells above the 99% quantile. Data was log-normalized. Samples were clustered individually by selecting the 3,000 highest variable genes with the FindVariableFeatures function from the *Seurat* package. From the overlap of these lists, the top 3,000 genes were selected to calculate principal components (PCs). Top 30 PC embeddings were adjusted with *harmony* R-package, with samples as covariates. In the resulting integrated feature space the nearest neighbor approach and graph based louvain algorithm implemented in *Seurat* was used to cluster cells and stepwise test optimal cluster resolution (from 0.1 to 1.6 in 0.1 steps) and computing silhouette widths. Celltype markers were calculated with the FindMarkers function with default parameters (wilcoxon test) in *Seurat* and cell types were manually annotated based on known canonical markers.

We removed four distinct small clusters that were inconclusive for different reasons, i.e. high expression of mitochondrial genes, expression of multiple cell type markers, consistently low RNA and Feature counts. After removal the integration process was repeated and a final atlas was created.

### Composition analysis

We tested if cell type or state composition changes between groups are meaningful by implementing a permutation approach to estimate a null distribution. For each individual cell, we considered the sample it came from and the cell type label it was assigned. From this table, we created 1000 permutations. For each permutation run we calculated the cell proportions for each sample and calculated the mean proportion per cell per group (control, HF), from which the difference in cell proportion was calculated as test statistic. By calculating the proportion per sample and not per group, we simulated unequal cell numbers in samples. The resulting 1000 random cell proportion differences are an estimate for a null distribution (Supp. Fig. 3A). All distributions passed Shapiro-Wilk test for normality (p>0.05). We calculated the area under the normal curve from the mean and standard deviation of the null distribution to estimate the probability of observing the actual measured proportional difference (Supp. Fig. 3B).

### Sample distance

To prioritize cell types displaying disease signatures we calculated distances between cell types per sample^42^. First, highly variable features were calculated per cell type with *FindVarFeature* function from *Seurat* and the top 1000 features were selected for distance calculation. For each cell type and sample, pseudobulk profiles were TMM normalized and voom transformed with the *edgeR* and *voom* R-package and cosine distances were calculated. We calculated median sample distances within groups and between groups to assess the distance ratio. Cell types with distance ratio below 1 show higher sample distances between groups than within groups and are candidates for differential gene expression analysis.

In addition to sample distance, we applied the R-package *Augur*^9^ to train random forests to classify the experimental group (control vs. heart failure) of individual cells. We used the *calculate_auc* function with default parameters.

### Differential gene expression analysis

To control for different absolute numbers of cells per sample we subsampled the total number of cells to the lowest cell number in a sample. For these cells we calculated differentially expressed genes with *FindMarker* function from *Seurat* R-package. To ameliorate sampling effects we repeated this subsampling process 5 times, and reported the gene intersection of genes with Benjamini Hochberg corrected p-value <0.05 and absolute log2FC >0.1. Upregulated genes were considered as the disease signature of the respective HF model.

### Cell state analysis

To identify cell states in macrophages, we subset each sample to macrophages and reintegrated samples by following the same steps as described above. For fibroblasts, the integrated atlas was used to calculate meaningful distances between cells via construction a nearest neighbor graph. Cell states were then defined by the louvain clustering algorithm implemented in *Seurat’s FindCluster* function, and optimal cluster resolution (0.1 to 1 in 0.1 steps) was determined by selecting the resolution with maximal silhouette width. Cellstate interpretation was aided by processing cell states reported in^19^ from steady and perturbed state cell markers.

### Functional Analysis

We performed functional analysis of top 100 cell state markers and fibrotic signatures. Overrepresentation analysis was performed with *enrichR,* with GO-molecular function and biological function terms. Additionally, functional gene sets were acquired from MSIG DB and subjected to hypergeometric testing. Pathway analysis was performed with PROGENy^83^. To calculate cell state pathway activities, we summed up cells per cell state to create pseudo bulk profiles, which were analyzed for pathway activities. For the integrated atlas we relied on running pathway analysis per cell to not sum uncorrected counts to pseudobulks. For study comparison we used log fold change as an effect size reported per study to calculate progeny scores. We used TF regulons obtained from *DoRoThEA*^84, 85^ and the *decoupleR*^84, 85^ R-package to estimate TF activities. We used univariate linear models to estimate TF activity on logFC vectors from different studies. Finally, we calculated module scores which are weighted expression means for genesets with the *AddModuleScore* function in *Seurat*.

### Study Integration

Two additional 10x Genomics scRNAseq datasets were analyzed by downloading raw FASTQ files and processing via cell ranger pipeline as described above. Sample integration was performed via canonical correlation analysis as implemented in *Seurat*. Unsupervised clustering and cluster marker assessment was used to identify fibroblasts in each study, which were subset to perform study integration. We integrated fibroblast cell data from three datasets via calculating highly variable features in each dataset, using 3000 overlapping features of all datasets. We used *Harmony* with study and sample ID as covariates for dataset integration. Downstream analysis was performed as described above. Integrated data was reclustered to identify cell states and markers for each cluster were calculated based on log transformed data. To evaluate integration performance we ensured that each study contributed cells to each cluster (Supp. Fig. 6). To quantify batch effects from different studies, samples and experimental groups, we calculated a batch mixing score based on average silhouette width as proposed previously^86^. A score of 1 represents a balanced integration while 0 represents strong batch effect conservation. The Integrated fibroblasts atlas yielded a batch score of ∼0.99 for study labels, ∼1 for group labels and ∼0.97 for sample labels. To avoid study batch effects in differential expression analysis, we calculated differentially expressed genes between control and disease models per study. We performed downsampling to equalize cell numbers per sample as described earlier. We collected genes that appeared in at least 4 of 5 downsampling runs by passing Benjamini Hochberg adjusted wilcoxon p-value <0.05.

### Assessing state dependency of transcriptional shifts

To estimate transcriptional shifts of disease signatures, we estimated how well these signatures separate healthy and disease cells within an assigned cell state. We first calculated gene set scores for each cell of the respective HF model via the *AddModuleScore* function from *Seurat* R-package (Supp. Fig. 6A). The difference of these gene set scores was then assessed by calculating the area under the receiver operator curve (AUROC) (Supp. Fig. 6B) as a metric for the transcriptional shift within a state.

For a single gene being expressed in a state dependent manner we expected that expression levels would vary between states. To quantify this dependency, we fit ANOVA models for each gene of the disease signatures by modeling their expression value by IFS category (gene X ∼ IFS) and extracted the explained variance of the model (eta² values). To compare these with disease related variance we fit ANOVA models for the same genes but with group labels (gene X ∼ group). The ANOVAs were calculated for all HF models separately.

### Cell-cell communication

We used two approaches to estimate cell-cell communication. First, we performed ligand-receptor (LR) analysis with the method aggregation tool *LIANA*^56^. Second, we used *NichNet*^57^ to connect ligands with receiver cell gene expression^57^. To lower the false positive rate of LR pairs, we only analyzed macrophages and fibroblasts as both cell types displayed the strongest disease response. We ran *LIANA* on control and HFpEF mice separately. Within each group we aggregated ranked LR pairs from 6 LR tools with the *aggregate_l* function in *LIANA*. We assumed that an LR pair is specific for HFpEF if it is ranked low in control and high in HFpEF mice. To quantify this we calculated a HFpEF specificity score (*S*) by: *S* = (1 − *RankCT*) * *RankHF*.

We selected top 30 LR pairs based on *S* and filtered for ligands being expressed by more than 0.15 percent in HFpEF sender cells and to be upregulated >0.1 log2FC; receptors were filtered for expression percentage >0.15 in HFpEF receiver cells (Supp. Fig. 9E). The top ligands from macrophages were used as input for *NicheNet* to model regulatory potential for the HFpEF disease signature. Since the prior knowledge graph in NicheNet is undirected, directional regulation of target genes cannot be modeled. For this reason we used up-and downregulated genes in HFpEF fibroblasts as the target cell gene set.

### Human Bulk RNA-seq

For human HFpEF bulk analysis, raw count data was downloaded from European nucleotide archive, accession number E-MTAB-7454. We filtered genes by a minimum of two RNA counts in at least 30% of samples per experimental group. We TMM normalized samples and voom transformed for variance stabilization and performed DEA with *limma* and *edgeR* R-packages. HFrEF bulk data from the *Reference of the Heart Failure Transcriptome* (ReHeaT)^44^ is available at: https://zenodo.org/record/3797044#.XsQPMy2B2u5. We selected the top 500 genes for overrepresentation analysis.

### Murine bulk RNA-seq of different timepoints from HFpEF hearts

Mice of the HFpEF model were used after 10 (n=4) and 15 (n=4) weeks of diet protocol and compared to mice that received control diet (n=3). After sacrificing the animals by cervical dislocation, the still beating heart was directly placed in ice-cold HBSS where atria and large vessels were dissected and immediately placed into liquid nitrogen and stored at −80°. The tissue was homogenized and total RNA isolated using TRIzol (Thermo Fischer Scientific). Quantification and quality controls were performed by DS-11 spectrophotometer (DeNovix) and 5300 Fragment Analyzer System (Agilent) and samples used with RNA Integrity Number (RIN)>8 (mean RIN 9.2 ± 0.5 SD). Library construction and sequencing was performed by the GeneCore Facility at EMBL Heidelberg. Briefly, libraries were prepared from 1 µg total RNA using respective Illumina mRNA Kits according to the manufacturer’s instructions. Samples were multiplexed and single-end sequencing performed on an NextSeq 500 (Illumina). Raw BCL data were demultiplexed and then converted to FASTQ files. Reads were aligned via ArchS4 pipeline implemented in BioJupies^87^. We filtered lowly expressed genes and normalized samples using the trimmed mean of M-values (*edgeR* ^88^) and subsequent variance-stabilizing transformation (limma voom) and performed differential expression analysis (*limma*^89^). Resulting t-values were used for enrichment analysis via run_ulm function from *decoupleR*^84, 85^ R-package.

### Flow cytometry

In order to obtain a single-cell suspension, the removed hearts were minced and digested in 450 U/mL collagenase I, 125 U/mL collagenase XI, 60 U/mL DNase I, and 60 U/mL hyaluronidase (MilliporeSigma) for 1 hour at 37°C under agitation. Cells were washed, counted and diluted to 100µl per 10×10^6^ cells prior staining. The whole cardiac sample was further proceeded for and passed through multiparameter flow cytometry analysis. An additional control sample was used for unstained and fluorescence-minus-one gating controls. Spleens were passed through a 40µm cell strainer and washed prior to further staining. Peritoneal macrophages were collected directly after sacrificing the animal by peritoneal lavage performed by injecting 5ml of ice-cold PBS with 3% FCS using a 27G needle. After an abdominal massage for 30sec the fluid was collected by a plastic pipette through a small incision of the abdominal wall. Cells were kept on ice, washed and resuspended for 1:100 antibody staining. For all samples, Fc receptor blocking was performed for 10min prior to fluorescent antibody staining using anti-CD45-PerCP-Cy5.5 (BD Biosciences, clone 30-F11), anti-CD11b-APC-Cy7 (BD Biosciences, clone M1/70), anti-Ly6C-BV605 (Biolegend, clone AL-21), anti-F4/80-PE-Cy7 (Biolegend, clone BM8), anti-MHCII-BV421 (BD Biosciences, clone M5/114.15.2), anti-CCR2-APC (R&D Systems, clone #475301). For lineage exclusion PE-conjugated anti-Ter119 (BD Biosciences, clone TER-119), anti-NKT (BD Biosciences, clone U5A2-13), anti-B220 (BD Biosciences, clone RA3-6B2), anti-CD49b (BD Biosciences, clone DX5), anti-90.2 (BD Biosciences, clone 53-2.1) and anti-Ly6G (BD Biosciences, clone 1A8) antibodies were used. Gating strategies and representative plots are presented in the supplementary figures (Supp. Fig. 11). Flow cytometry was performed on a FACSCelesta (BD) and data analysis conducted by FlowJo (BD) software.

### Histological analysis

Harvested hearts were rinsed in PBS, and fixed for 7 days in 10% buffered formalin at room temperature. Hearts were subsequently dehydrated, paraffinized, and sectioned (5 μm). Cardiac fibrosis was assessed by using the cardiac muscle Picrosirius-Red Stain Kit (abcam, ab245887) according to the manufacturer’s instructions. Immunohistochemistry (IHC) was performed using heat-mediated antigen retrieval in sodium-citrate buffer at pH 6.0 and an anti-rabbit secondary antibody containing HRP/DAB Detection IHC Kit (abcam, ab64261) according to the manufacturer’s instructions. The following primary antibodies were used: anti-ANGPTL4 (#710186, ThermoFisher Scientific, clone 1HCLC), anti-CILP (CAU24345, Biomatik), anti-FAP (ab207178, Abcam). Slides were mounted and imaged using a brightfield (SP2, Leica) and fluorescence microscope (Axio Observer, Zeiss).

### Immunofluorescence

For immunofluorescence staining, in TissueTek embedded and frozen heart-derived cryostat sections were fixed in 3,7% PFA, permeabilized for 20min, blocked with 5% BSA for 1h, and stained overnight with anti-CollagenIV (1:200, ab6586, Abcam) and the following day for one hour with goat anti-rabbit AlexaFluor594 (1:400, ab150080, abcam) secondary antibody. Nuclei were stained with a DAPI-containing mounting medium, covered and images captured using a Axio Observer (Zeiss) fluorescence microscope.

### FAPI PET-CT

We performed murine PET-CT imaging with 68Ga-FAPI-46 as described previously^90^. Briefly, the radiotracer (ca. 13MBq), produced and approved for patient application, was injected via the tail vein of isoflurane anesthetized mice and 10min static images recorded using a small-animal PET/CT scanner (Inveon; Siemens).

### HFpEF patient population

The study protocol is in accordance with the declaration of Helsinki and has been approved by the local ethics committee of the University Hospital Heidelberg and was registered on ClinicalTrials.gov (Identifier Number: NCT04317911). Written informed consent was obtained prior to participation. Details, inclusion criteria, patient characteristics and further methods were described previously^59^. Briefly, 102 patients, admitted for atrial fibrillation cryoablation having an EF > 50%, were enrolled prospectively and screened for HFpEF by echocardiography, stress-echocardiography, 6-minute-walking-test and blood biomarker tests. 20 patients fulfilling the current HFpEF diagnosis criteria of the Heart Failure Association (HFA) of the ESC^91^ were compared with 20 propensity score matched controls derived from the enrolled non-HFpEF atrial fibrillation patients. Clinical follow-up was performed after 6 and 12 months including baseline examinations and 24h holter ECG.

### Human ANGPTL4 plasma ELISA

Peripheral venous blood was drawn from the right femoral veins during vessel access prior to percutaneous pulmonary vein isolation and plasma aliquots stored at −80°C. A human ANGPTL4 solid-phase sandwich ELISA kit (#EHANGPTL4, ThermoFisher) was used according to the provided manufacturer’s protocol. Samples were analyzed in doublets, 100 µL 1:2 diluted plasma incubated overnight under agitation at 4°C. Absorbance was detected at 450 nm using an EnSpire (PerkinElmer Inc) plate reader. The standard curve and final protein concentrations were calculated using Prism 9 (GraphPad). ROUT test for outlier identification was performed (Q=0.1%) and one outlier in the HFpEF group was excluded, which exhibited a lipemic plasma sample.

## Acknowledgements

Immuno-Fib HF supported by a grant from the Leducq Foundation (20CVD02). Research grant from the German Society of Cardiology to LMW (No. DGK15/2020). FL is supported by the Heisenberg Programm of the German Research Foundation (DFG). JDL, RORF, JHS and JSR are supported by Informatics for Life funded by the Klaus Tschira Foundation. We acknowledge the great technical support by Dr Tina Fuchs and Dr. Volker Ast of the NGS core facility for single-cell genomics at Universitätsmedizin Mannheim. Biorender was used for illustration of figure schematics. We thank Susann Pohl for excellent technical assistance and Jiale Huang for help with the animal model. We acknowledge Leonie Küchenhoff for providing critical feedback that improved the manuscript.

## Author Contributions

F.L, L.M.W. and J.D.L. conceived and designed the study. L.M.W. carried out the experiments and analyzed the data. N.H. supported the histological analysis. M.M.Z. collected patient plasma samples and clinical parameters. J.D.L. curated data, conceived and performed formal analysis. R.O.R.F and J.S.R advised data analysis. J.D.L and L.M.W. wrote the original draft of the manuscript. F.L., J.S.R., R.O.R.F, N.H. and F.S. reviewed and edited the manuscript. F.L., J.S.R., J.H.S. and N.F. supervised the project and acquired funding.

## Data and Code availability

Code that is used in the analysis is accessible under https://github.com/saezlab/scell_hfpef. Single-cell and bulk RNAseq data of murine HFpEF model will be published in the Gene Expression Omnibus after manuscript publishing. Other datasets used in this study are available online through referenced publications (MI study, E-MTAB-7895; AngII study, E-MTAB-8810).

## Supplementary Tables

Table 1. Integrated fibroblast cell state marker

Table 2. Fibroblast disease signatures by study

