## Supplemental_Material for "Single-cell transcriptomics reveal distinctive patterns of fibroblast activation in murine heart failure with preserved ejection fraction"

#shared last authorship

[1] Institute for Computational Biomedicine, Heidelberg University, Heidelberg, Germany

[2] Faculty of Biosciences, Heidelberg University, Heidelberg, Germany

[3] Internal Medicine II, Heidelberg University Hospital, Heidelberg, Germany

[4] Informatics for Life, Heidelberg, Germany

[5] Department of Cardiology, Internal Medicine III, Heidelberg University Hospital, Heidelberg, Germany

[6] German Center for Cardiovascular Research (DZHK), Partner Site Heidelberg, Heidelberg, Germany

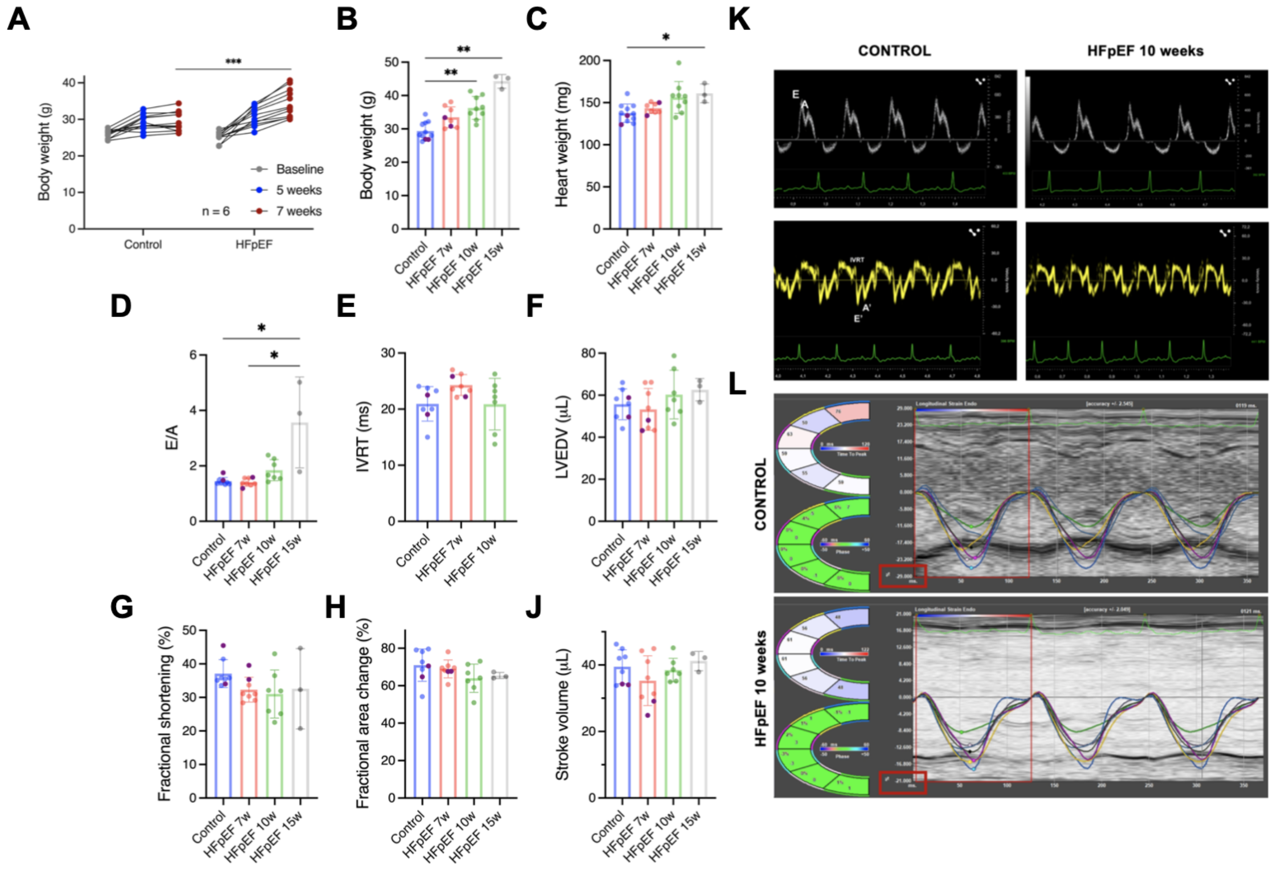

**Supplemental Figure 1. HFpEF disease model characterization.**

A) Body weight increase in mice between diet start and 7 weeks diet showing a significant increase in body weight after 7, but not after 5 weeks. B) Body weight of all animals at the time point of final echocardiography. C) Heart weight after organ removal. D) Ratio between early diastolic LV blood flow over the mitral valve to late diastolic filling induced by atrial contraction. E) Isovolumic relaxation time (IVRT) measured in tissue doppler traces. Bad ultrasound conditions of obese 15w mice prevented robust IVRT measurements. F) Left ventricular end-diastolic volume (LVEDV) derived from semi-automated LV-traces of the parasternal long axis. G) Myocardial fractional shortening in M-mode traces. H) Fractional area change measured by semi-automated traces from the parasternal short axis. J) Cardiac stroke volume, acquired as volumes in F. K) Representative pulse wave doppler (upper panels) and tissue doppler (lower panels) images. L) Example image of global longitudinal strain in the parasternal long axis analyzed by VevoStrain. A-J display mean ± SD, statistical analysis performed by Students t-test (A) or one-way ANOVA, *$p<0.05$, **$p<0.01$, ***$p<0.001$.

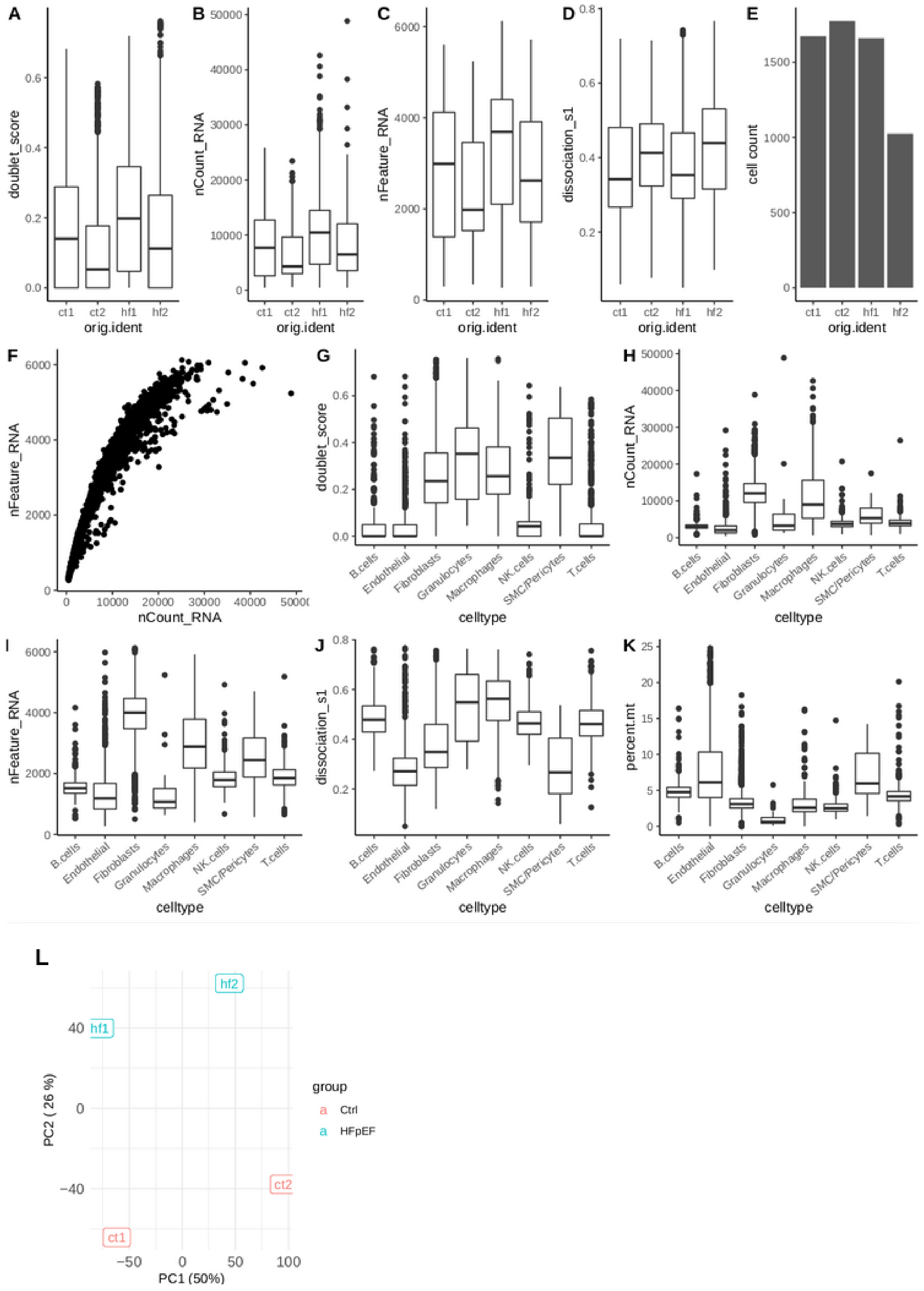


**Supplemental Figure 2. Quality control of single cell RNAseq data.**

A) Doublet score distribution per sample calculated with scDoubletFinder.B) Total RNA count per sample. Mean of all cells is 8,240. C) Total feature count per sample (Mean of all cells

2,838). D) Dissociation score (weighted mean of dissociation associated gene set) computed per sample. E) Total cell count per sample. F) RNA and feature count for all cells after filtering (< 6000 & > 300 features). G) Doublet score distribution across cell types. H) RNA count distribution across cell types. I) Feature count distribution across cell types. J) Dissociation score distribution across cell types. K) Mitochondrial percentage distribution across cell types. Boxplots in A-D, G-K display median and IQR. L ) Principial component analysis of sample-level pseudobulks. A batch effect associated with sequencing run is associated with PC1, while PC2 associates with the contrast of diet.

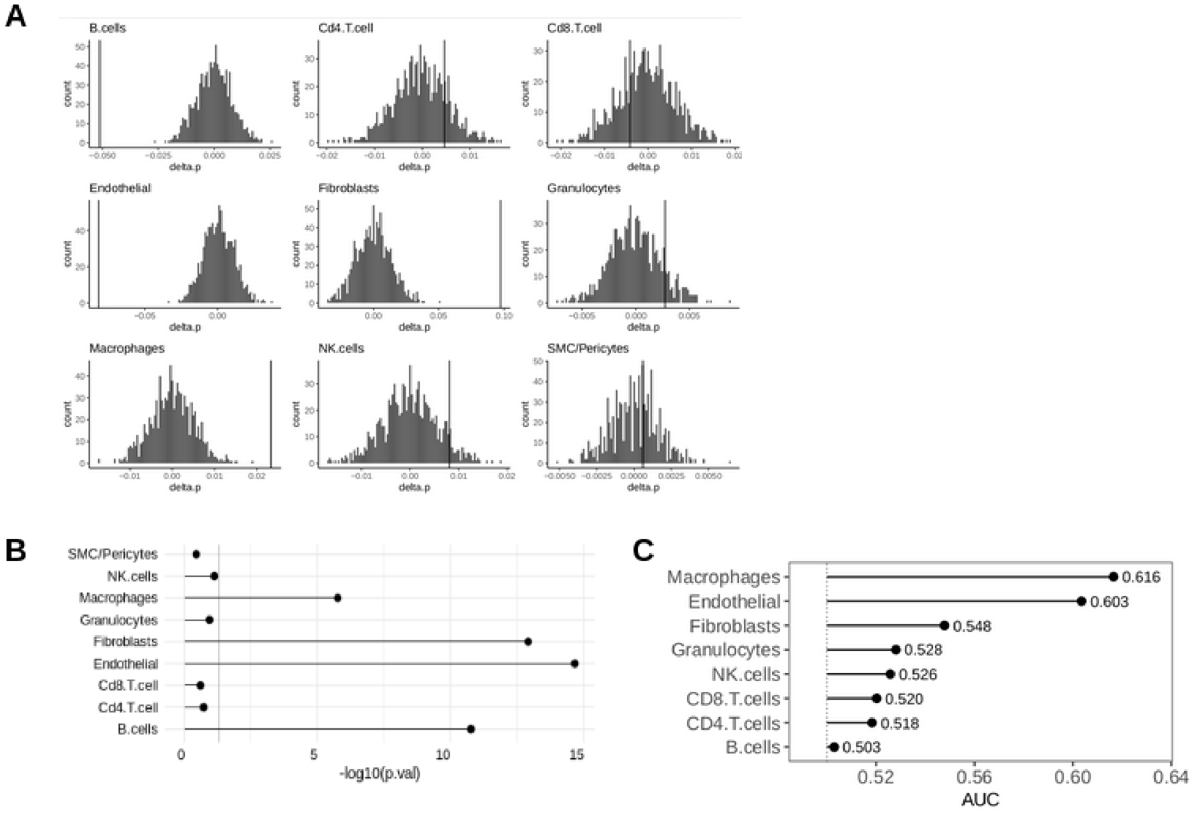


**Supplemental Figure 3. Label Permutation for composition analysis.**

A) Histograms of compositional difference (delta.p) generated by label randomization. Vertical lines indicate observed difference. B) P-value of compositional differences based on normal distribution inferred from A) per cell type. Vertical line at 0.05. C) Classification success by cell type measured by the area under the receiver operator curve (AUC) based on the *Augur* method.

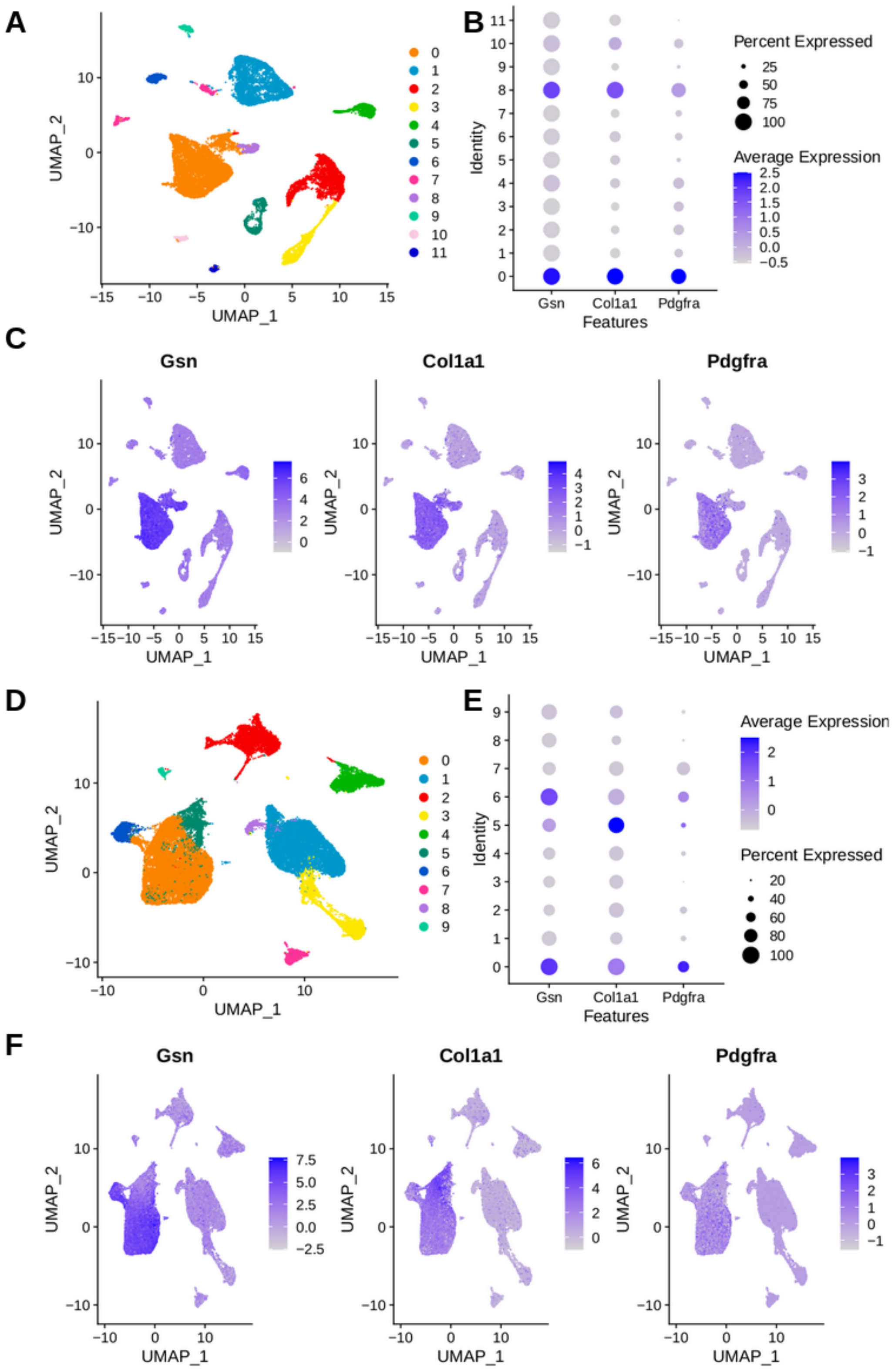


**Supplemental Figure 4. Fibroblast annotation in AngII and MI model.**

AngII model (A-C) and MI model (D-E). A, D) UMAP embedding of full cell atlas after processing, filtering and clustering. B) Expression of fibroblast marker genes (Gsn, Col1a1, Pdgfra) in AngII model per cluster. Cluster 0 and 8 were subset for downstream analysis. E) Expression of fibroblast marker genes (Gsn, Col1a1, Pdgfra) in MI model per cluster. Cluster

0, 5 and 6 were subset for downstream analysis. C, F) Expression of marker gene UMAP.

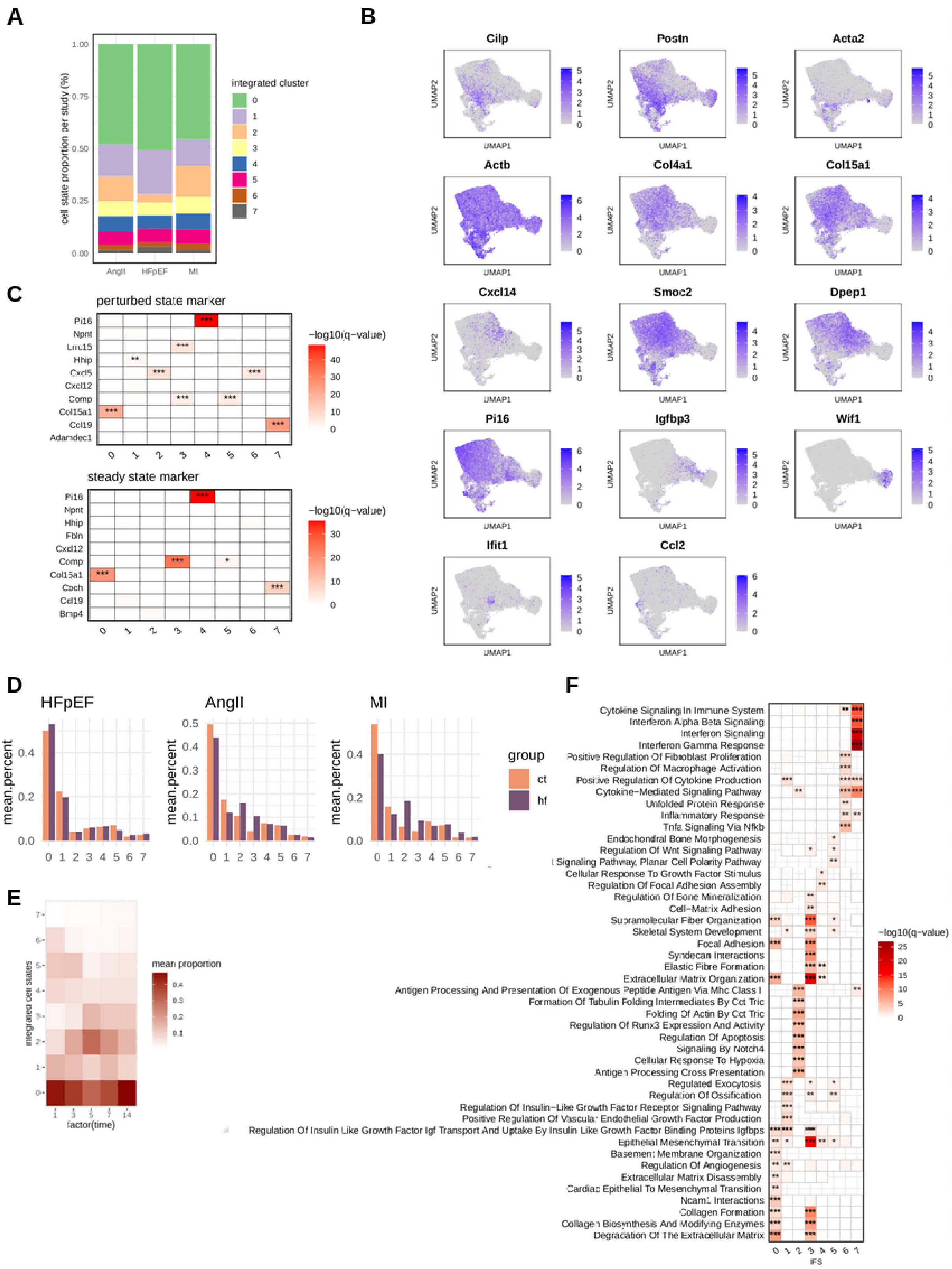


**Supplemental Figure 5. Integrated cell state (IFS) comparison.**

A) Composition of integrated cell states per study. B) UMAPs colored by selected gene marker expression across the integrated fibroblast atlas. C) Overrepresentation analysis of

IFS markers with markers derived from a perturbed and steady state cross-organ fibroblast atlas. D) IFS composition separated by experimental group (color) and HF models (panels). E) IFS composition of the myocardial infarction samples by time points in days (x-axis). F) Overrepresentation analysis of MSIG DB gene sets with IFS state marker. C+F, hypergeometric test, with Benjamini Hochberg correction, *q < 0.05, **q < 0.01, ***q < 0.001.

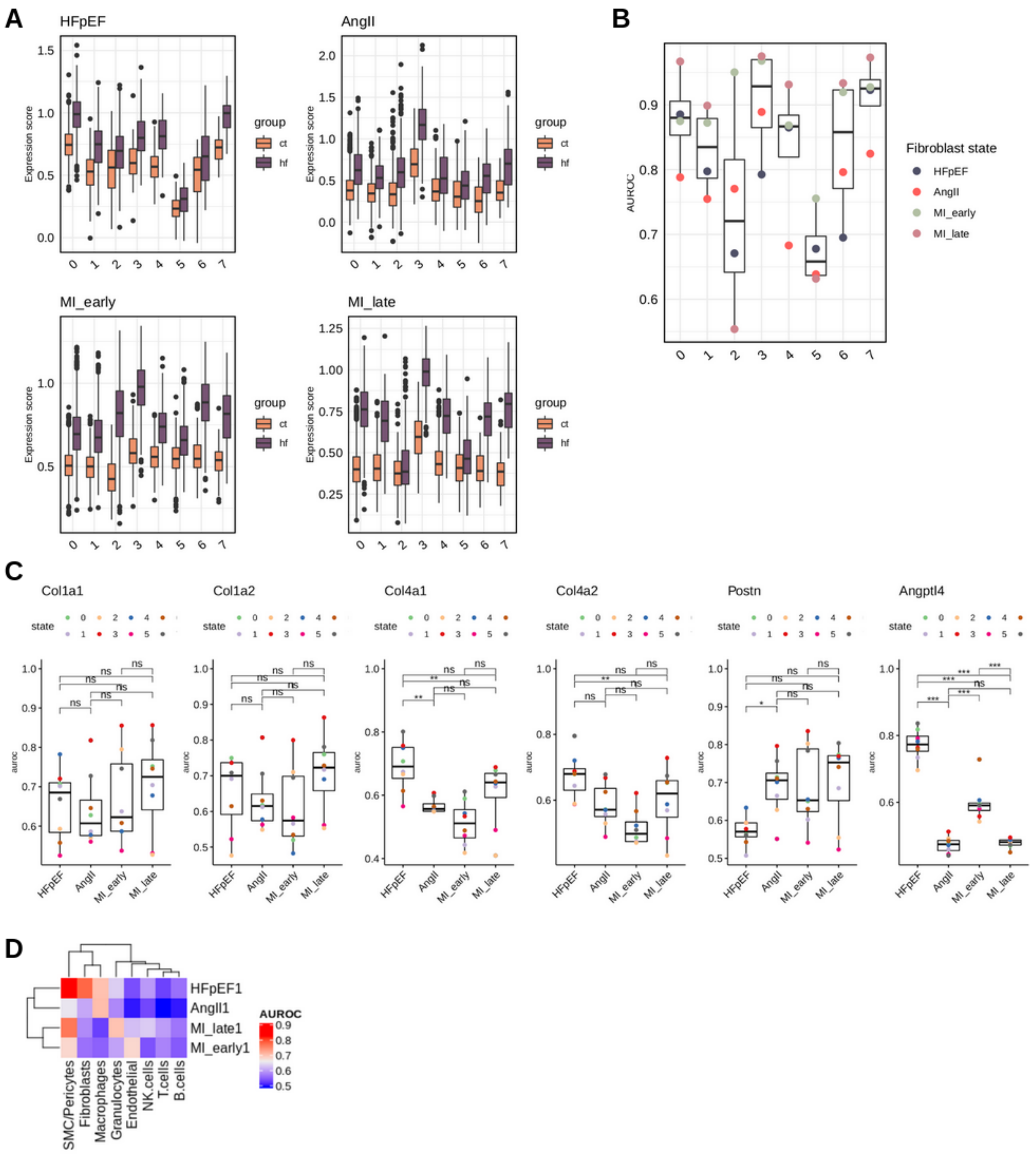


**Supplemental Figure 6. Assessing transcriptomic shifts across mouse models.**

A) Gene set scores of disease signatures calculated for individual cells, across IFS (x-axis), disease groups (color) and studies (panels). B) Quantification of IFS separability by module scores from panel A, assessed via area under the receiver operator curve (AUROC). C) Quantification of IFS separability assessed via AUROC based on expression of selected genes, Col1a1, Col1a2, Col4a1, Col4a2, Postn and Angtpl4 per heart failure model. Wilcoxon test, *p < 0.05, ** p < 0.01, *** p < 0.001.

D) Assessing possible background expression. Fibroblast signatures (y-axis) are summed to module scores per cell and used to calculate AUROCs between control and heart failure samples within the HFpEF data set for each cell type (x-axis).

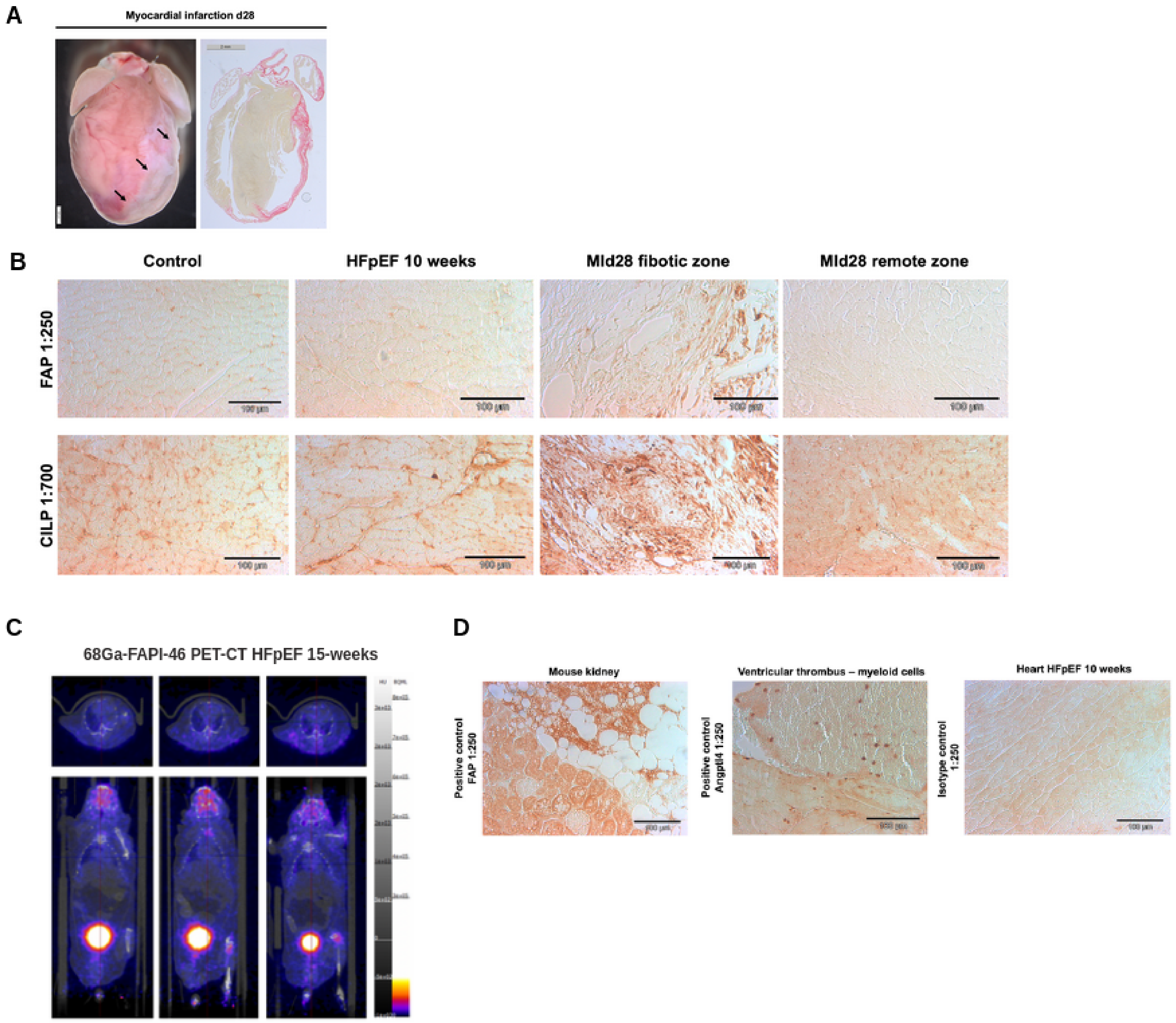


**Supplemental Figure 7. CILP and FAP protein expression in HFpEF and MI mouse models.**

A) Demonstration of the large infarct area 28 days after minimal-invasively induced myocardial infarction. B) Immunohistochemistry DAB stainings of FAP and Cilp in control, HFpEF 10 week-diet hearts and of the fibrotic and remote zone 28 days after myocardial infarction (MI). C) Images of 68Ga-FAPI-46 PET-CT in three individual HFpEF mice after 15 weeks of dietary intervention. Upper panels show transverse and lower panels show coronal planes of the heart. D) Immunohistochemistry validation of FAP using mouse kidney tissue for positive control as recommended by the manufacturer. Validation of Angptl4 staining by intracardiac blood leukocytes as positive control. Right panel shows a respective image of isotype control stainings.

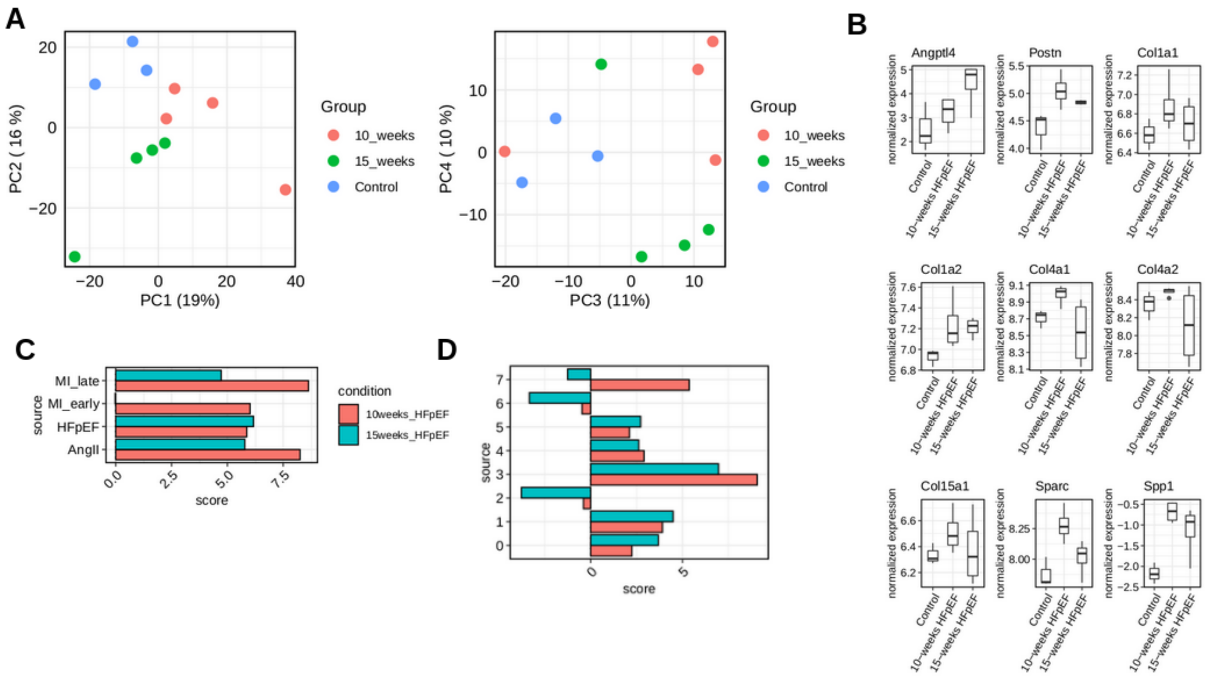

**Supplemental Figure 8. Corroborating fibroblast disease signature**

A) Principal component analysis of bulk transcriptomics of mice; three control, four 10-week and four 15-week of treatment with L-NAME and high fat diet. B) Single gene expression (TMM normalized and log transformed) of selected genes. No gene reached significance. C+D) Enrichment analysis of different gene sets (y-axis) in t-values from contrast analysis (15-week vs control, green; 10-week vs control, red). Score represents univariate linear model parameters. C) Enrichment of fibroblast disease signatures from different murine heart failure models. All scores were significant ($p<0.05$), except early MI signature in 15 weeks of HFpEF. D) Enrichment of integrated fibroblast state markers (y-axis). IFS 0, 1 ,2, 3, 4, 5 enrichment scores were significant in both contrasts;  IFS 7 enrichment score was significant in 10 weeks only ($p<0.05$).

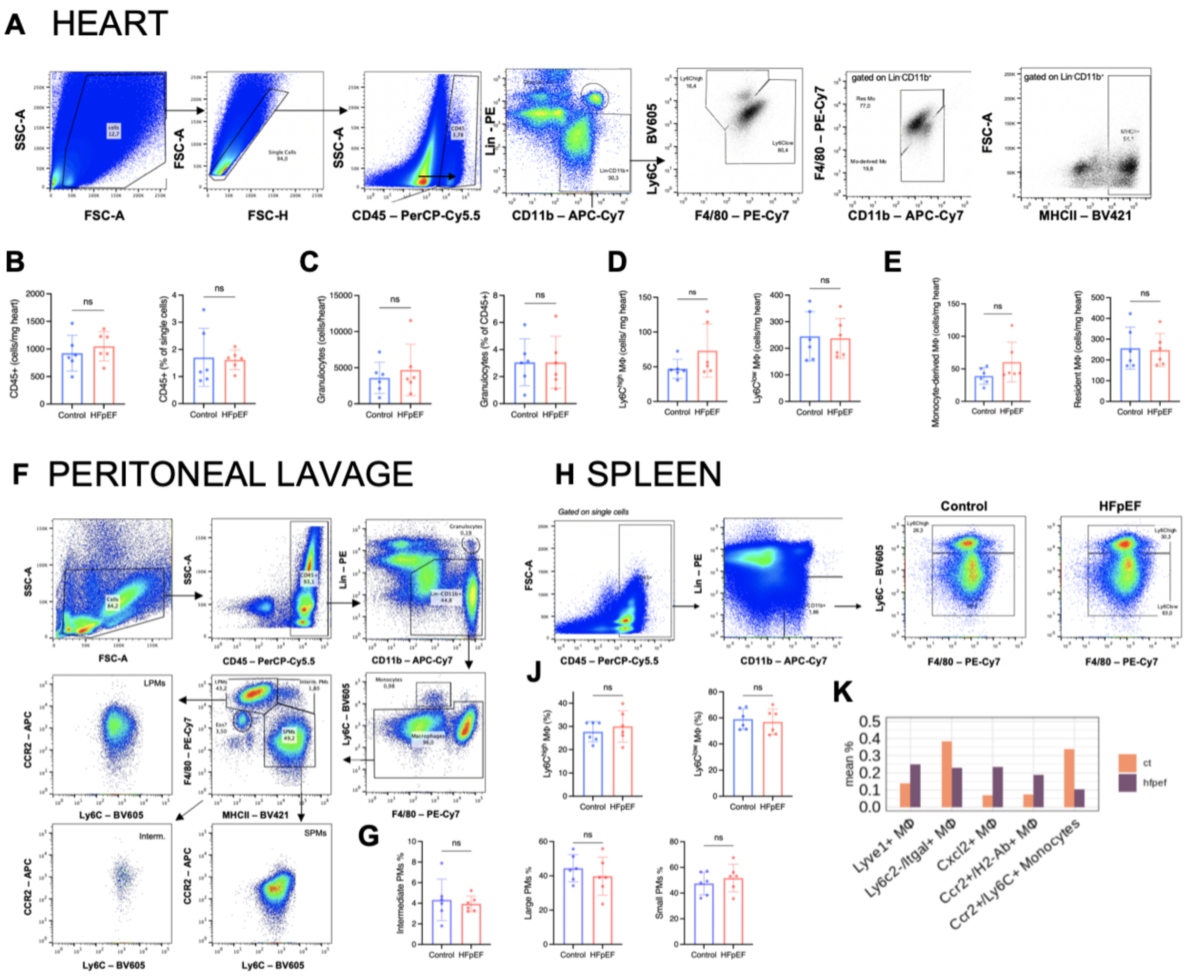


**Supplemental Figure 9. Flow cytometry of macrophages (MΦ) in HFpEF.**

A) Flow cytometry gating strategy of heart samples. B-E) Quantification of absolute leukocyte flow cytometry results, obtained by running complete organ samples through the cytometer. F) Flow cytometry gating strategy of peritoneal macrophages derived from peritoneal lavages. LPMs, large peritoneal macrophages; Interm, intermediate macrophages; SPMs, small peritoneal macrophages. G+J) Quantification of relative peritoneal (G) and spleen (H) monocytes/ macrophages (MΦ). H) Flow cytometry gating strategy of spleen MΦ and representative plots of Ly6C$^{high/low}$ monocytes/ macrophages (MΦ). Statistical analysis using unpaired t-test, n=6/group, bar graphs indicate mean±SD, ns= not significant. K) Macrophage cellstate composition (mean percentage per group) comparison between control and HFpEF, from scRNAseq. No statistical analysis performed, due to low cell counts.

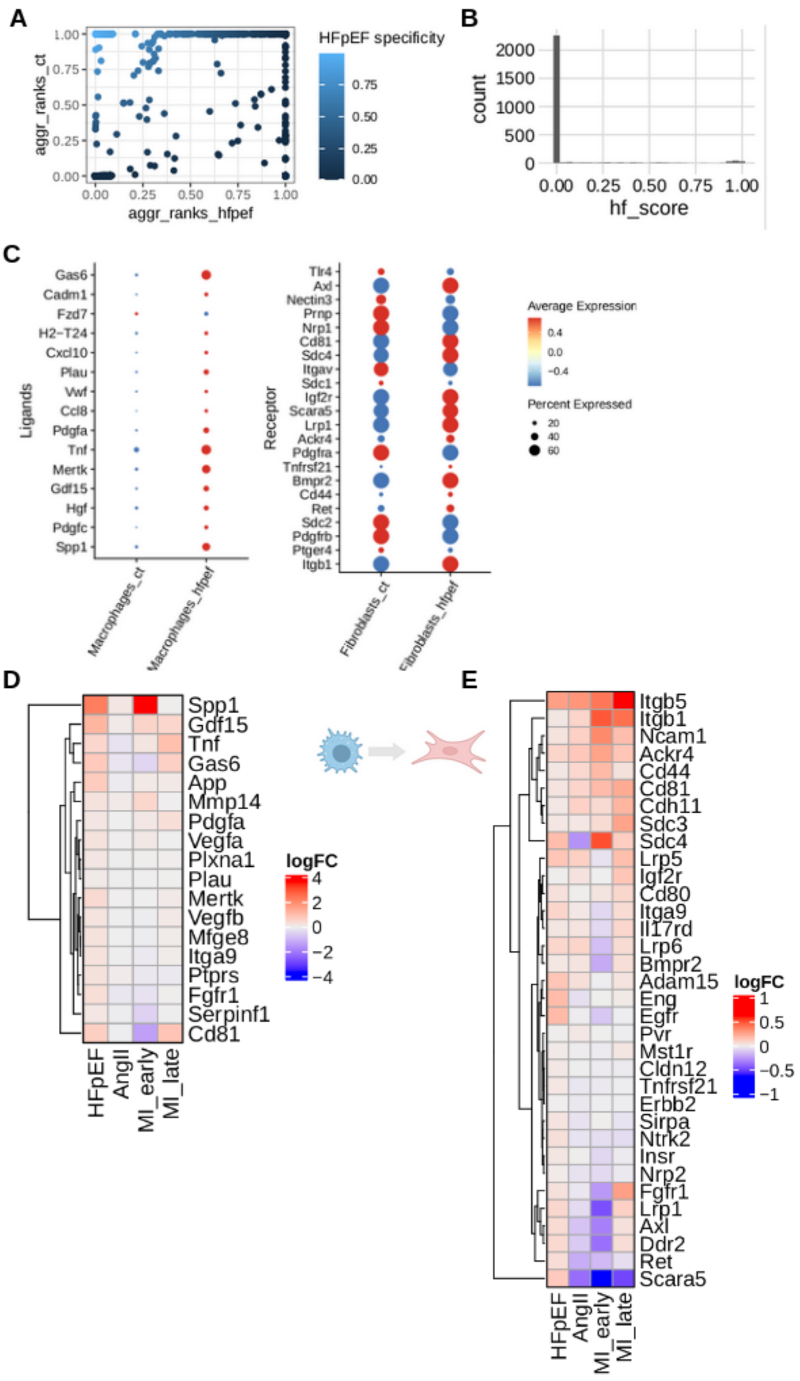


**Supplemental Figure 10. Ranking of Ligand Receptor (LR) pairs between Macrophages and Fibroblasts.**

A) Aggregated ranks from HFpEF (x-axis) and control (y-axis) mice, coloring represents HF score (see methods). LR pairs in the upper left corner received high ranks in HFpEF mice and low ranks in control mice. B) Distribution of the HFpEF specificity score (hf_score). We selected LR pairs with hf_score > 0.9 for downstream analysis. C) Selected LR pairs and their expression between control and HFpEF mice. Ligands (left panel ) and receptors (right panel) expressed in Macrophages and Fibroblasts, respectively. D+E) Comparison of log fold change regulation in heart failure of ligands in macrophages (D) and receptors in fibroblasts (E) across heart failure models.
